# A quantitative risk assessment approach for longline fishing gear impacts on seafloor habitats

**DOI:** 10.1101/2025.09.15.676354

**Authors:** Beau Doherty, Lisa Lacko, Allen R. Kronlund, Sean P. Cox

## Abstract

Bottom longline fishing gear used worldwide to capture fish and invertebrate species can impact seafloor habitats, leading to increased use of spatial closures (e.g., MPAs) in areas where habitat risks are considered high. However, such closures often rely on limited data, because fishing impacts on habitat are rarely quantified and fine-scale habitat maps are often unavailable. In this paper we develop fine-scale species distribution models for coral and sponge habitats and demonstrate a quantitative risk assessment framework for habitat impacts from bottom longline trap and hook fisheries, using the British Columbia Sablefish fishery as a case study. We estimate a 4% (95%CI: 2-7%) reduction in coastwide sponge habitats due to Sablefish fishing from 1965 to 2024, compared to pre-fishery levels. Habitat status at finer spatial scales of 1 km^2^ shows similar trends to the coastwide aggregate status, with habitat declines less than 10% for 99% of fishing grounds. Our analysis provides fine-scale information on habitat distribution and the impacts from Sablefish longline trap and hook fishing gear, providing key information for conservation planning and fisheries management. Our risk assessment approach provides quantitative metrics (relative benthic status) for ecosystem objectives focused on fishery impacts on habitat. Such habitat metrics can be incorporated into fisheries management strategy evaluation, allowing resource managers to compare performance of alternative strategies against a broader suite of sustainability objectives that include habitat, fish stocks, and fisheries catch.

## 1. Introduction

Bottom-contact fishing gears such as trawls, longlines, and traps are used worldwide to capture fish and invertebrate species that inhabit the seafloor. Bottom fisheries pose a risk to sensitive benthic habitats such as corals or sponges that are considered Vulnerable Marine Ecosystems (VMEs, Auster et al., 2011) due to their susceptibility to disturbance (Krieger and Wing, 2002; Heifetz et al., 2009) and can take decades to recover from damage (Andrews et al., 2002, 2009). Spatial information on fishing risks and habitat is essential for effective VME management that balances conservation goals and economic benefits for fisheries. Without this information, poorly placed closures risk shifting fishing effort into more vulnerable habitats outside protected areas, while also limiting access to economically valuable fishing grounds where habitat risks are low. Effective spatial management decisions for fisheries and VMEs requires the use of quantitative habitat assessment metrics that account for cumulative fishing impacts.

VMEs are considered biodiversity hotspots that provide essential habitat for deep-sea ecosystems and are prioritized in global marine conservation planning (Auster et al., 2011; Kaikkonen et al., 2024). Bottom-contact fisheries are perceived as a primary threat to VMEs, which has led to increasing implementation of spatial closures such as Marine Protected Areas (MPAs) to restrict fishing in areas where risks are perceived as high. Yet such management strategies for protecting VMEs are often based on limited data because risks from bottom contact fisheries are poorly understood (Pitcher et al., 2016; Kaikkonen et al., 2024) and fine-scale mapping for VMEs may not be available.

Understanding the risks that fishing poses to deep-sea habitats and VMEs is critical for managing bottom-contact fisheries impacts on ecosystems, yet quantitative frameworks for assessing habitat status and fishing impacts are rarely used in fisheries management and conservation planning (Welsford et al., 2014a; Pitcher et al., 2016, 2022; Kaikkonen et al., 2024). Due to data limitations for habitat mapping and assessing fishing impacts, qualitative frameworks for habitat risk assessment (Hobday et al., 2011) are more commonly used (Kaikkonen et al., 2024; Hobday et al., 2011). These non-quantitative frameworks are typically based on subjective rankings of habitat susceptibility to damage from fishing gears and expectations for recovery rates (i.e., resilience) (Eno et al., 2013; Williams et al., 2011), which are prone to bias because they rely heavily on analyst assumptions or expert opinions (Hordyk and Carruthers, 2018).

Such qualitative frameworks provide a relative ranking of habitat risk that can be useful for prioritizing fishing areas that require more detailed risk assessment approaches. For example, the hierarchical framework for the Ecological Risk Assessment for the Effects of Fishing (ERAEF, Hobday et al., 2011), allows qualitative or semi-quantitative approaches to be used for lower risk activities, while higher risk activities undergo quantitative risk assessment. Such quantitative metrics are essential for evaluating alternative management strategies that consider trade-offs between conservation goals and economic objectives (Pitcher et al., 2016). Just as stock assessments and management strategy evaluation (MSE) are used to quantify the stock status of target species to evaluate such trade-offs, similar frameworks could assess habitat status by accounting for fisheries’ cumulative impacts and modelling habitat population dynamics.

In this paper, we demonstrate a quantitative risk assessment framework for bottom habitats that is broadly applicable to longline fisheries, here focused on the Sablefish (*Anoplopoma fimbria*) fishery in British Columbia (BC), Canada. The main requirements are a habitat map, estimates of fishery bottom contact, and recovery rate information for sensitive benthic taxa. This approach was first developed as part of the Canadian Sablefish Association’s (CSA) Research Program for risk assessment and management strategy evaluation of Sablefish fishing impacts on the SGaan-Kinghlas - Bowie Seamount Marine Protected Area (Doherty et al., 2018b; Rossi et al., 2018). Following more data collection and improved estimates of longline trap and hook bottom contact for the fishery (Doherty et al., 2025), we can now apply this approach to the coastal Sablefish fishery in BC, providing a case study for a quantitative risk assessment approach for longline fisheries bottom impacts. We measure risk to sponge habitats within Sablefish fishing grounds by estimating current habitat status relative to pre-fishery (i.e., unfished or pristine) levels via logistic population growth models for discrete 1 km^2^ spatial units, similar to the relative benthic status (RBS) approach previously applied for trawl fisheries (Pitcher et al., 2016, 2022). Assessments of sponge habitat status are then available for each spatial unit and the aggregate coastwide area where the fishery operates. To demonstrate this approach, we consider only Sablefish fishing impacts on habitat status, while future assessments could consider cumulative impacts of other bottom fisheries that overlap with Sablefish fishing grounds in BC. The main steps in our risk assessment include “*(i) an integrated system of data capture technology for Sensitive Benthic Habitats (SBHs) containing corals and sponges, (ii) in situ monitoring and estimation methods to quantify bottom contact from fishing gear, (iii) presence–absence* [and density] *modelling to map SBHs, (iv) risk assessment for quantifying bottom fishing impacts on SBHs and spatial assessment of conservation status, and (v) a simulation framework for evaluating fishing impacts on SBHs and their recovery trajectories under alternative management approaches*” (Doherty et al., 2025). Steps (i) and (ii) have been previously described by (Doherty et al., 2025). This paper focuses on (iii) and (iv) by developing fine-scale maps of presence-absence and density for coral and sponge habitats and a spatial logistic population model for habitat that accounts for annual impacts from historical Sablefish fishing effort and subsequent recovery. Step (v) will be completed as part of future research initiatives for the BC Sablefish bottom contact research program.

## 2. Methods

Here we described the methods for implementing the first 4 steps (i-iv) of the quantitative risk assessment approach described above (Doherty et al., 2025) for assessing risks to SBHs from bottom longline trap and hook gear deployed in the BC Sablefish Fishery from 1965-2024. First, we describe the deep-water cameras and sensors (Step 1) deployed on fishing gear to collect presence-absence observations of habitat and measure gear movement (Doherty et al., 2018a). Second, we quantify habitat-gear contact (Step 2) from historical Sablefish longline trap and hook gear (step ii, Doherty et al., 2025). Third, we explain habitat modelling (Step 3) to produce fine-scale predictive maps of coral and sponge density and distribution (step iii, Doherty et al., 2021). Fourth, we describe the spatial risk assessment (Step 4) of sponge habitat, using inputs from steps i and ii, which includes mortality from gear contact and recovery via a logistic population model to assess habitat status (Pitcher et al., 2016, 2022).

### 2.1. Step 1. Data Collection

The CSA and Fisheries and Oceans Canada (DFO) have collaborated on bottom contact research since 2013 (Doherty et al., 2018a, 2021). Deep-water cameras, accelerometers, and depth sensors were deployed on Sablefish longline trap gear in BC to collect observations of deep-sea habitats and quantify habitat-gear contact during fishing operations (Doherty and Cox, 2017; Doherty et al., 2018a, 2021). In-situ observations of bottom habitat were collected from trap-cameras deployed on the annual stratified random survey (StRS) for BC Sablefish from 2013–2017 (Wyeth et al., 2007). Trap cameras were mounted inside individual traps deployed on longline sets, typically with 26 total traps. Cameras were programmed to record 1-minute videos within stationary traps on the seafloor or when accelerometers triggered video recordings during gear movement (Doherty et al., 2018a).

All epifauna observations from camera video were identified to the lowest possible taxonomic rank, typically to the Order or Family level. Taxa identification from underwater video is frequently limited due to restricted visibility and the absence of physical samples, the latter of which is often needed to identify sponges and corals to genus or species level (Williams, 2013; Austin et al., 2013). More information on taxonomic identification and images from trap-camera are provided in Doherty and Cox (2017) and Doherty et al. (2018a).

### 2.2. Step 2. Habitat-gear contact from Sablefish fishing

Here we described the methods used to estimate habitat-gear contact from Sablefish longline trap and hook fisheries from 1965–2024, using the same approach previously used to estimate the Sablefish footprint for 2007–2023 (Doherty et al., 2025). We use the term footprint to describe gear-contact area with the substrate and seafloor habitats, the latter of which can occur for gear interactions with taller habitat structures slightly above the seafloor (Doherty et al., 2025).

The following sections describe methods for generating i) fisheries effort time series from 1965–2024 (2.2.1), ii) longline trap gear footprints (2.2.2), iii) longline hook gear footprints (2.2.3), and iv) combined annual footprints for both longline hook and trap gear (2.2.4). We provide only brief summaries of methods used for ii) and iii) here, as they have been previously described in detail by Doherty et al. (2025),

#### 2.2.1. Sablefish fishing effort

We use commercial fishery effort data from 1990–2024 sourced from fisher logbooks stored in the Fisheries Operations System (April 2006 to December 2024) and PacHarvSable (January 1990 to March 2006) databases at the DFO. We include records from trips targeting Sablefish and combination trips targeting both Pacific Halibut (*Hippoglossus stenolepis*) and Sablefish in our analysis, the latter of which was first permitted under the BC groundfish management plan in 2006. We use ‘Sablefish fishing’ throughout the paper to encompass both directed Sablefish trips and combined Pacific Halibut and Sablefish trips using either longline hook or trap gear.

Sablefish fishing records are well documented with reliable set coordinates since the introduction of the individual vessel quota system in 1990 (Turris, 2000), but earlier records are largely incomplete. Prior to 1990 there are coastwide catch data but limited information on the spatial coordinates of sets or total effort. For the 1965–1989 period we estimate annual traps and hooks deployed by the Sablefish fishery using catch and commercial catch-per unit effort (CPUE) data for longline trap gear (*g* = 1) and longline hook gear (*g* = 2), using the steps described below.

For commercial longline trap (LLT) fisheries, we use standardized commercial trap CPUE (kg/trap) that is available from 1979–2009 and used as an abundance index in Sablefish operating models (Johnson et al., 2024). We extend this time series back to 1965 using estimates of catchability 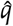 and vulnerable biomass *B* for the trap fishery (*g* = 1) from the Sablefish operating model (OM, Johnson et al., 2024)

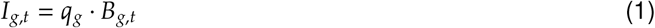

where *I*_*g,t*_ is an estimated CPUE index for *g* = 1 and year *t* ∈ {1965, 1966, …, 1978}.

For commercial longline hook (LLH) fisheries, we use annual CPUE indices (kg/hook) generated by dividing the longline gear Sablefish catch by the total number of hooks deployed in each year across all Sablefish fishing trips

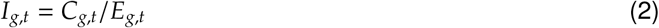

where *I*_*g,t*_ is an estimated CPUE index *g* = 2 for longline hook gear and *t* ∈ {1990, 1991, …, 2024}. We then use CPUE *I*_*g,t*_ and vulnerable biomass *B*_*g,t*_ for the longline hook fleet *g* = 2 (Johnson et al., 2024) to calculate annual catchability *q*_*g,t*_ using the same approach as equation (1), solving for *q*.

We then calculate 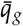 using the mean annual log catchability log *q*_*g,t*_ from the last 5 years of complete data

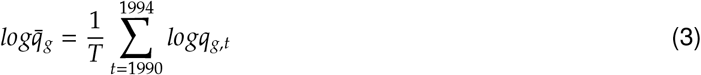

where *T* = 5 years, covering the period from 1990 to 1994. We adopt 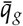 as the catchability estimate from 1984–1989; however, we apply an adjustment factor for the 1965–1983 period to account for the switch from j-hooks to circle hooks in 1984. The widespread adoption of circle hooks across the entire fleet is estimated to have increased CPUE by a factor of 2.2 (Williams and McCaughran, 1985; IPHC, 1985). Longline hook CPUE index series for 1983–1989 and 1965–1983 are obtained by multiplying 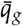 and 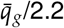, respectively, by the vulnerable biomass for longline hook gear (1).

Annual effort *E*_*g,t*_ is then calculated as total number of traps (*g* = 1) or hooks (*g* = 2) deployed for each gear type *g* in year *t* by dividing catch *C*_*g,t*_ by CPUE *I*_*g,t*_ (i.e., solving for effort in 2),

For longline trap fisheries (*g* = 1), *I*_*g,t*_ is an estimated index for *t* ∈ {1965, 1966, …, 1978} and *I*_*g,t*_ via (1) and is observed data for *t* ∈ {1979, 1980, …, 1989}, while *I*_*g,t*_ is an estimated index via (1) for *t* ∈ {1965, 1966, …, 1989} for longline hook fisheries (*g* = 2).

Trap fisheries typically deploy 60 traps per longline set (Doherty et al., 2025; Johnson et al., In Review), which we use to convert units of historical trap effort *E*_*g,t*_ to total sets for longline trap deployments from 1965–1989. Similarly, we convert total hooks effort into sets using 955 hooks per set, which is the mean number of hooks deployed per longline set from Sablefish and Halibut/Sablefish combination trips from 2010–2021 (Johnson et al., In Review).

We allocate longline trap and hook sets from 1965–1989 by random sampling with replacement from the historical set distribution from 1990–2024 for each gear type, weighted according to the proportion of total sets in each grid cell. To account for uncertainty, we repeat this sampling to produce *n* = 100 replicates of annual spatial effort distribution from 1965 to 1989 for each gear type.

#### 2.2.2. Longline trap gear contact

We estimate bottom footprints from Sablefish longline trap gear 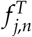 by multiplying estimates of the trap footprint length *l* by the footprint width *w* for trap *n* on set *j*

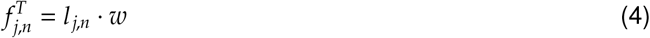

We use an estimate of *w* = 96 cm for sponge taxa for a conical Sablefish trap with 137 cm (54 inch) bottom hoop diameter (Doherty et al., 2025).

Footprint lengths are estimated using a model that accounts for the effects of fishing depth *D*_*j*_ and the trap retrieval position *X*_*j*_, *n* (e.g., first, middle, last) on gear movement (Doherty et al., 2025)

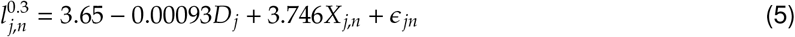

Parameters for eqn. 5 are estimated from a linear model fit to outputs from a movement classification algorithm, which converts video, depth, and accelerometer observations from sensors deployed on trap gear into estimates of trap drag lengths (Doherty et al., 2025). The *ϵ*_*jn*_ term is an independent and identically distributed normal residual (i.e., 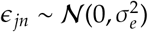) where *σ*_*e*_ = 1.2.

The trap retrieval position *X*_*n*_ is calculated relative to the total number of traps *N* deployed on the set as:

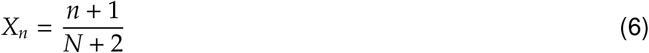

This allows us to account for sets with different numbers of traps deployed by expressing the trap retrieval position as a percentage of overall set length. We add 1 to numerator and 2 to the denominator to account for anchors at both ends of the set (Doherty et al., 2025). When depth estimates were not available in fishing records, we extract the depth at the set midpoint from GEBCO bathymetry data (GEBCO Compilation Group, 2024).

#### 2.2.3. Longline hook gear contact

We estimate longline hook gear footprints 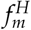 for set *m* by multiplying the set length *l* by the lateral line movement *w*

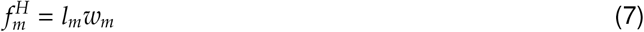

The set lengths *l* are calculated assuming a straight line between start and end deployment locations. In cases where set lengths are unavailable due to missing deployment locations, we use the median set length of 3.1 km from Sablefish longline sets from 1990–2024. We use estimates of *w* = 6.2 m (95% CI: 2.5-9.9) from the Patagonian Tootfish fishery (Ewing et al., 2014; Welsford et al., 2014b), which uses similar gear.

#### 2.2.4. Annual habitat-contact from fishing gear

For the 1965-1989 period where spatial coordinates of fisheries effort are unavailable, we calculate the footprint (e.g., gear-habitat contact area) *f*_*c,g,t*_ in each 4 km x 4 km grid cell *c* for gear type *g* and time *t* by multiplying the number of sets *N*_*c,g,t*_ by the mean footprint per set 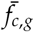 by grid cell and gear type derived from the 1990–2024 LLT and LLH footprint estimates (described in sections above). The annual footprint *f*_*c,t*_ for each grid cell *c* for *t* ∈ {1965, 1966, …, 1989} is then calculated by summing footprints across both gear types:

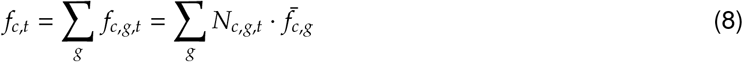

For the more recent period from 1990 to 2024 with reliable records of set coordinates, we sum the total habitat contact area *f*_*c,t*_ from LLT and LLH sets to estimate the cumulative footprint for trap sets (*j*=1,2,…,*J*_*c,t*_) and hook sets (*m* = 1,2,…, *M*_*c,t*_) in each grid cell *c* for year *t* ∈ {1990, 1991, …, 2024}

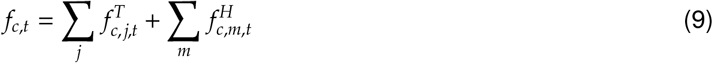

We then calculate the proportion of the grid cell *λ*_*c,t*_, assuming no overlap of footprints from each set as

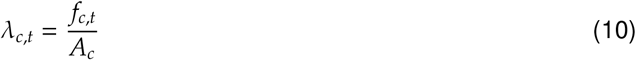

where *A*_*c*_ = 16 km^2^ for 4 km x 4 km grid cells.

To account for potential overlap of sets contacting habitat, we assume that sets are randomly distributed within each grid cell *c* in year *t* according to a Poisson distribution (Gerritsen et al., 2013; Amoroso et al., 2018), using the probability mass function

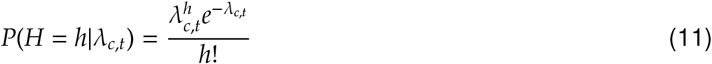

where *H* is a discrete random variable with parameter *λ* > 0 and *h* is the number of times that habitat is contacted. We then estimate the proportion of habitat in each cell *c* contacted at least once (i.e., *h* ≥ 1) as 1 minus the probability of no habitat contact.

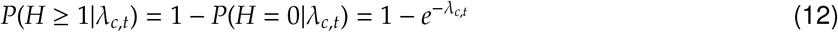

### 2.3. Step 3. Habitat Distribution Modelling

#### 2.3.1. Predictor variables for habitat modelling

We identified 8 predictor variables for model fitting as potential indicators of sponge and coral habitats (Whitney et al., 2005; Leys, 2013; Winship et al., 2020; Gonzalez-Mirelis et al., 2021). Variables included depth, slope, broad-scale bathymetric position index (BPI), a hard bottom index, temperature, aragonite saturation rate Ω_*arag*_, bottom current, and primary productivity. All variables were converted to raster layers with 100 m × 100 m resolution for predicting sponge and coral habitats within the BC Sablefish fishery footprint (Doherty et al., 2025) and survey area (Wyeth et al., 2007), primarily within depths ranging from 200-1400 m, using the terra package in R (Hijmans, 2024; R Core Team, 2024). We used the Albers Equal Area projection (EPSG: 3005) for BC with a spatial extent of 546 km × 746 km (longitudinal range: 134.0^°^ to 125.7^°^ W, latitudinal range: 48.1^°^ to 54.6^°^ N).

Depth for the majority of the prediction grid was obtained from a 10 m resolution digital elevation model (DEM) for BC (Kung, 2023). We merged the 10 m DEM (Kung, 2023) with additional 5 m resolution multibeam survey data from Queen Charlotte Sound and the west coast of Graham Island (unpublished data, R Kung, Natural Resources Canada), as well as a 3-arc second DEM (Carignan et al., 2013) to extend depth data into deeper waters in our prediction grid off of Haida Gwaii and Queen Charlotte Sound. Slope was derived from the depth raster using the 4 neighbours method (Fleming and Hoffer, 1979) with the terra package in R (Hijmans, 2024). We calculated a broad-scale BPI (1 km inner radius, 10 km outer radius) using the MultiscaleDTM (Lundblad et al., 2006; Ilich et al., 2023) package in R.

We obtained seasonal mean values, averaged from the 1981–2010 period, for bottom temperature, bottom aragonite saturation rate, bottom current, and primary productivity from the BC continental margin model (BCCM, Masson and Fine, 2012; Peña et al., 2019) at a 3km spatial resolution. We used mean values across all seasons and resampling with bilinear interpolation to generate a 100 m resolution for the prediction raster grid. We also considered BCCM bottom layer data for alkalinity, dissolved inorganic carbon, oxygen, and salinity as predictor variables; however, these were all strongly correlated (Pearson correlation coefficients of 0.92-1) with aragonite and temperature. To avoid collinearity, we fit models with either aragonite and temperature, but not both, using aragonite for coral models and temperature for sponge models.

Bottom substrate classifications (rock, mixed, sand, or mud) for BC at 100 m spatial resolution were obtained from a random forest model (Gregr et al., 2021). We generated a hard bottom index with a continuous scale of 0 to 1 for each 100 m grid cell by calculating the proportion of neighbouring cells within a 10-cell radius (1 km) that were classified as rock or mixed bottom types (Nephin et al., 2023) using the terra package in R (Hijmans, 2024; R Core Team, 2024).

#### 2.3.2. Trap bottom location estimator

We use a Bayesian trap location estimator (Doherty et al., 2018a, 2021) to account for differences between the surface deployment locations and the location of fishing gear on the bottom. This generates posterior probabilities for potential trap landing locations where habitat observations occur. We used the trap-camera surface deployment locations as uncorrelated bivariate normal prior distributions for trap bottom longitude *x* and latitude *y* for *k* sets (Doherty et al., 2021), estimated as:

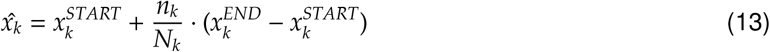

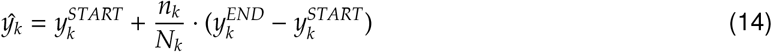

where *n* is the trap number deployed with cameras and *N* is the total number of traps deployed on the set. The *START* and *END* superscripts indicate latitude and longitude locations at the start and end of the set, respectively.

The posterior is proportional to the product of the surface deployment location prior and a likelihood function that uses the depth sensor observations at the bottom location and 10 m bathymetry data (Kung, 2023; Carignan et al., 2013), described above (For a description of the statistical model, see Table 3 in Doherty et al., 2018a). We renormalize the 10 m x 10 m grid cells to assign zero probability for cells with probabilities for trap landing locations less than 0.01% (Doherty et al., 2018a, 2021). The posterior probability grid from the Bayesian trap location estimator is used to weight the extraction of predictor variable rasters at each coral and sponge observation site, accounting for uncertainty in the trap bottom location (Doherty et al., 2021).

#### 2.3.3. Habitat distribution model

We develop separate spatial models for probability of sponge (Porifera) and coral (Alcyonacea) presence *p* and density *D* using generalized linear mixed effects models (GLMMs) with spatial random effects. Spatial random effects use a stochastic partial differential equation (SPDE) approach for linking Gaussian fields with Gaussian Markov random fields (Lindgren et al., 2011). We fit models with the sdmTMB package in R (Anderson et al., 2025; R Core Team, 2024), which uses Template Model Builder (TMB, Kristensen et al. (2016)) for model fitting and the fmesher package (Lindgren, 2023) to construct matrices for SPDE inputs.

We use a two-part delta modelling approach (Aitchison, 1955) with different processes for modelling presence-absence data and positive count observations. The presence-absence model uses a binomial distribution with a logit link function to model the probability of presence, while the positive count data are modelled via a Gamma distribution with a log link function, such that:

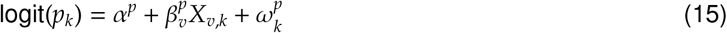

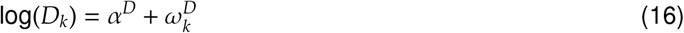

where *p*_*k*_ and *D*_*k*_ are the probability of presence and density, respectively, for observation *k, α* are the intercept terms, *β*_*v*_ are vectors of coefficients for *X*_*v*_ predictor variables, and *ω*_*k*_ is a spatial random field with a shared multivariate normal distribution *ω*_*k*_ ∼ MVN(0, Σ_*ω*_) for both *p*_*k*_ and *D*_*k*_ components. The *p* and *D* superscripts for *α* and *β* denote different coefficients for presence-absence and density models, respectively. The covariance matrices Σ_*ω*_ are constrained by a Matérn covariance function to define the rate of decline in spatial covariance with distance (Whittle, 1954; Matérn, 1986). We also tested initial density models that included environmental predictors; however, these models produced predictor responses that were either weak or ecologically implausible, suggesting potentially spurious relationships. Therefore, we modelled density using only an intercept term with spatial random effects, rather than attempting to model environmental effects on density with limited presence observations (n=16 for corals, n=28 for sponges).

For the SPDE approach we create a spatial mesh using a minimum distance of 5 km between knots with a correlation barrier so that spatial correlations declined 5 times faster over land than water (Bakka et al., 2019; Nephin et al., 2023). We use a penalized complexity (PC) prior on the Matérn parameters to constrain spatial random fields where *P*(*σ > σ*_0_) = 0.05 and *P*(*ρ < ρ*_0_) = 0.05, where *σ* is the spatial marginal standard deviation with a prior of *σ*_0_ and *ρ* is the distance (i.e. range parameter) at which two points are considered independent with spatial correlation of approximately 0.13 with a prior of *ρ*_0_ (Fuglstad et al., 2019). The choice for the *ρ*_0_ prior is expected to be greater than the true range with 95% probability and the choice for the *σ*_0_ prior is expected to be less than the true standard deviation with 95% probability (Anderson et al., 2025) based on simulation results that found: “*good coverage of the equal-tailed 95% credible intervals when the prior satisfies P*(*σ > σ*_0_) = 0.05 *and P*(*ρ < ρ*_0_) = 0.05, *where σ*_0_ *is between 2.5 to 40 times the true marginal standard deviation and ρ*_0_ *is between 1/10 and 1/2.5 of the true range*” (Fuglstad et al., 2019). We use priors of *ρ*_0_ = 13 km for coral models and *ρ*_0_ = 8 km for sponge models, which are 1/2.5th the initial range estimates fit without priors. For the spatial marginal standard deviation, we apply priors of *σ*_0_ = 42 and *σ*_0_ = 16 for corals and sponges, respectively, which are 20 times initial *σ* estimates fit without priors. We estimate shared parameters for range and standard deviation between the presence-absence and density models for model parsimony and since limited density observations did not allow for estimating density-model specific parameters.

The models are fit to the presence-absence *p* and density *D* observations from trap-camera videos of sponges (Phylum Porifera) and corals (Family Alcyonacea). Values for the depth predictor variable are measured in-situ during camera deployments by temperature-pressure recorders (Sea-Bird SBE 39), and when not available are extracted from bathymetry rasters based on surface drop location. Values for all other predictor variables were extracted from raster layers at observation locations, weighted by the posterior grid of probability for the trap bottom location. Prior to model fitting, predictor variables were centered and scaled by subtracting the mean and dividing by the standard deviation.

We initially included penalized smooth terms (Wood, 2017) for the depth covariate, which was thought to have potential for a quadratic relationship with habitat quality; however initial fits for presence-absence models showed weak depth effects without strong support for a non-linear effect. We subsequently fit the model with linear terms for all predictor variables. Initial model fits for sponge models indicated the hard bottom index had little effect, and therefore it was removed from subsequent fits. For corals, we excluded predictors for current and productivity after initial model fitting revealed they had little effect on presence-absence and did not improve model performance.

#### 2.3.4. Model Validation

We use 5-fold cross validation with spatial blocking (Roberts et al., 2017) to estimate AUC and *κ* statistics for evaluating model performance. To account for spatial autocorrelation, we divided the study area into spatial hexagons whose centres are separated by 30 km, which is 1.1 and 2.1 times the estimated autocorrelation range for presence-absence observations of corals and sponges, respectively, beyond which observations are considered independent. The observations within each hexagon are then randomly assigned to one of 5 folds using the blockCV package in R (Valavi et al., 2018), which uses an iterative approach to balance the number of presence and absence observations in each fold. For each fold, we hold out observations within that fold for testing and use the remaining 4 folds for fitting presence-absence models. We then calculate AUC and *κ* statistics for each fold.

We also assess goodness of fit by inspecting randomized quantile residuals (Dunn and Smyth, 1996) and simulation-based randomized quantile residuals (Waagepetersen, 2006), the latter of which uses the DHARMA (Hartig, 2022) package in R.

#### 2.3.5. Predicting habitat distribution

We generate habitat predictions for *i* ∈ {1, 2, …, 25, 081} 1 km x 1 km grid cells using predictor rasters described above, which were aggregated to 1 km resolution according to the mean values. We estimate standard errors and 95% confidences intervals in predictions by simulating uncertainty in fixed and random effects via 1000 draws from the joint precision matrix using the predict function in the sdmTMB package in R (Anderson et al., 2025).

#### 2.3.6. Coral and sponge habitat overlap within MPAs

We estimate coral and sponge habitat areas within existing and proposed MPAs in BC Sablefish fishing grounds following a five-step process:

1. Identify existing closures for bottom longline hook and trap fisheries within the Gwaii Haanas National Marine Conservation Area Reserve and Haida Heritage Site (GH NMCAR, Council of the Haida Nation and Parks Canada, 2018)that overlap with modelled habitat area (i.e., the Sablefish fishery and survey footprint)
2. Identify proposed MPAs within the Northern Shelf bioregion (NSB) with coral or sponge conservation objectives (Appendix 1 in MPA Network BC Northern Shelf Initiative, 2023) that overlap with modelled habitat area
3. Calculate area of overlap between modelled habitat and each MPA identified in steps 1) and 2) and exclude any closures with less than 5% overlap
4. Estimate the expected coral and sponge habitat in each grid cell by multiplying the grid cell area by their predicted probability of presence
5. Using the habitat estimate from 4), estimate the coral or sponge habitat area within closures

### 2.4. Step 4. Benthic Habitat Risk Assessment

In this section we use habitat-contact estimates from Sablefish fishing gear (2.2) with density models for sponge habitats (2.3), both described above, as data inputs for spatial risk assessment for sponge habitats. We use a logistic population model in each *i* 1 km x 1 km grid cell to estimate current habitat size in numbers *N*_*t*_ for each year *t* from 1965 to 2024

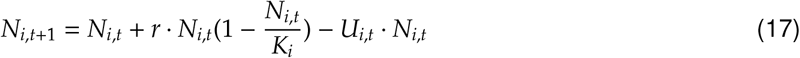

where *N*_*i,t*_ are numbers in each grid cell *i* at the start of year *t, U*_*i,t*_ is the annual sponge mortality rate in each grid cell, *r* is the intrinsic growth rate of the population, and *K* is the sponge habitat carrying capacity in the absence of Sablefish fishing. We assume that habitat was at carrying capacity prior (i.e., unfished habitat levels) to Sablefish fishing in 1965, as is typical in stock assessment, and exclude impacts from other bottom fisheries that fall outside the scope of the current assessment. We estimate the annual mortality rate in each grid cell *U*_*i,t*_ as the proportion of habitat contacted at least once within a year by Sablefish longline trap and hook gear (eqn. 12). This estimate assumes 100% mortality for gear-habitat contact.

We run the population model for *n*=100 replicates in each grid cell to account for uncertainty in historical footprints, the intrinsic growth rate *r*, and habitat density models. We sample *n*=100 replicates for historical distribution of longline trap and hook sets from 1965–1985 (described in Step 2). For intrinsic growth rate *r*, we draw *n*=100 values from a beta distribution with *β*(15.0, 125.2) parameterized to have mean *r*=0.107 with an upper 95% CI of 0.163 based on estimates from Rooper et al. (2011). The model is initialized as unfished *K*_*i*_ in each grid cell in 1965 (*t*=1) by drawing *n*=100 density *D*_0_, *i* values from the 1000 draws of predicted density distibutions in each grid cell (described above in *Predicting habitat distribution* section). We convert density to unfished numbers *N*_0,*i*_ in year *t*=1 by assuming a fixed field of view *θ* for each camera drop

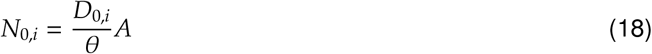

where *N*_0,*i*_ is equal to carying capacity *K*_*i*_ and *A* is 1 km^2^ for 1 km x 1 km grid cells. The *θ* field of view parameter is an unknown constant, which cancels out in eqn. 17. Although the field of view may differ across sets, for example depending on visibility and gear movement, the assumption of a constant field of view for each set is comparable to the assumption of constant search areas per unit of effort typically used for fisheries longline and trawl survey data.

We use this model (eqn 18) as our index of relative benthic status (RBS, Pitcher et al., 2016, 2022) for sponge habitats in 1 km x 1 km grid cells by dividing annual numbers *N*_*i,t*_ by carrying capacity *K*_*i*_ for each grid cell *i*

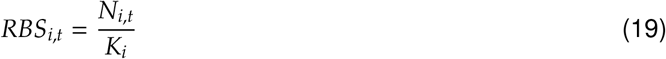

We calculate RBS for aggregated sponge habitats within BC Sablefish fishing grounds from 1965-2024 by summing the numbers *N*_*i*_ in each cell for time *t* and dividing by the summed carrying capacity *K* across all *i* grid cells

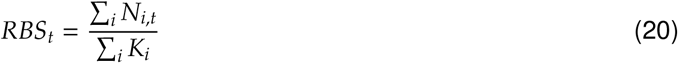

## 3. Results

### 3.1. Coral and Sponge Observations

Sponges were more frequently observed than corals with 28 presence and 254 absence observations (Fig. 1). The most commonly observed sponge taxa was *Auletta* sp., which is a smaller sponge ranging from 5–13 cm in height (Austin et al., 2013) that was observed on 11 trap camera deployments with high densities in some areas of up to 21 individual stalks (Table 1). Other sponge observations could only be identified to the class (Hexactinellida, Demospongiae) or phylum levels (Porifera), as is commonly the case for underwater video observations of sponges that often require physical samples to confirm species-level identification. Most locations had low sponge density with 1–3 counts per video, however a few locations in northwestern and western Haida Gwaii, and in Queen Charlotte Sound had higher densities ranging from 8–21 counts per video (Fig. 2).

**Table 1:** Summary of presence and density (mean and range of counts per trap camera deployment) for coral (Alcyonacea) and sponge (Porifera) taxa observed from trap cameras deployed in the BC Sablefish stratified random survey from 2013-2017

| Type | Lowest taxon identified | Locations observed | Density (Counts) | Depth range (m) |
| --- | --- | --- | --- | --- |
| Corals |  |  |  |  |
|  | Alcyonacea | 11 | 3.3 (1-25) | 552-1331 |
|  | Alcyoniidae | 1 | 1.5 | 1026 |
|  | *Heteropolypus* sp. | 1 | 1 | 552 |
|  | *Isidella* sp. | 2 | 1 (1-1) | 553-613 |
|  | *Paragorgia* sp. | 3 | 2.3 (1-3) | 220-687 |
|  | *Parastenella* sp. | 1 | 5 | 1265 |
|  | Primnoidae | 1 | 1 | 352 |
|  | *Swiftia simplex* | 2 | 2 (1-3) | 1026-1115 |
| Sponges |  |  |  |  |
|  | *Auletta* sp. | 11 | 6.8 (1-21) | 202-568 |
|  | Demospongiae | 8 | 2.2 (1-4) | 201-804 |
|  | *Farrea* sp. | 1 | 1 | 1265 |
|  | Hexactinellida | 5 | 1 (1-1) | 206-1331 |
|  | Porifera | 11 | 1.7 (1-3.5) | 245-687 |

**Figure 1:**
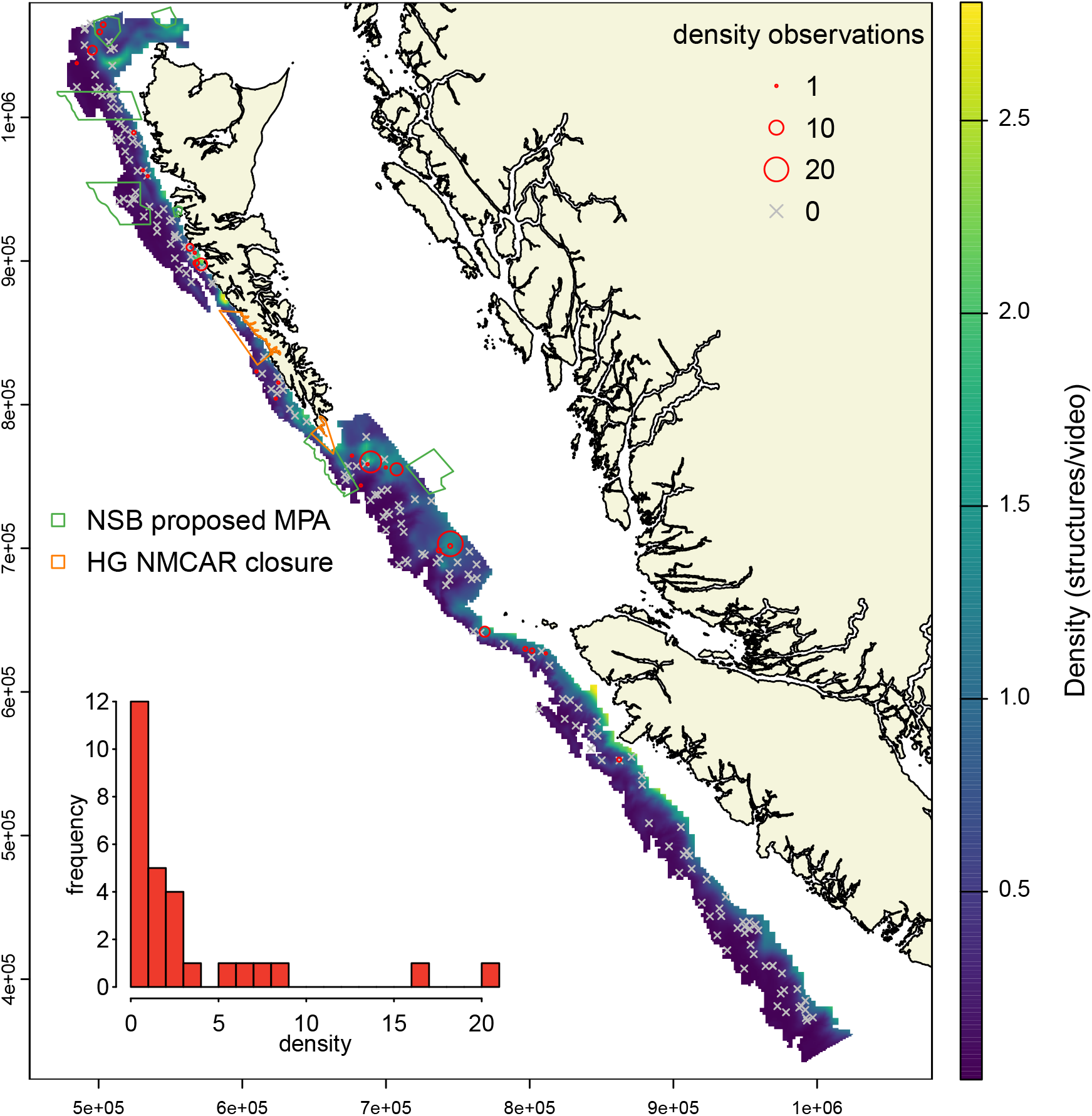
Mean predicted density for sponges from spatial models within Sablefish fishing areas in BC. Density (counts per video) for presence observations are shown as red circles sized in proportion to density, while absence observations are shown as grey Xs. Coastline data are from Wessel and Smith 1996 and map projection is NAD83 BC Albers (EPSG:3005).

**Figure 2:**
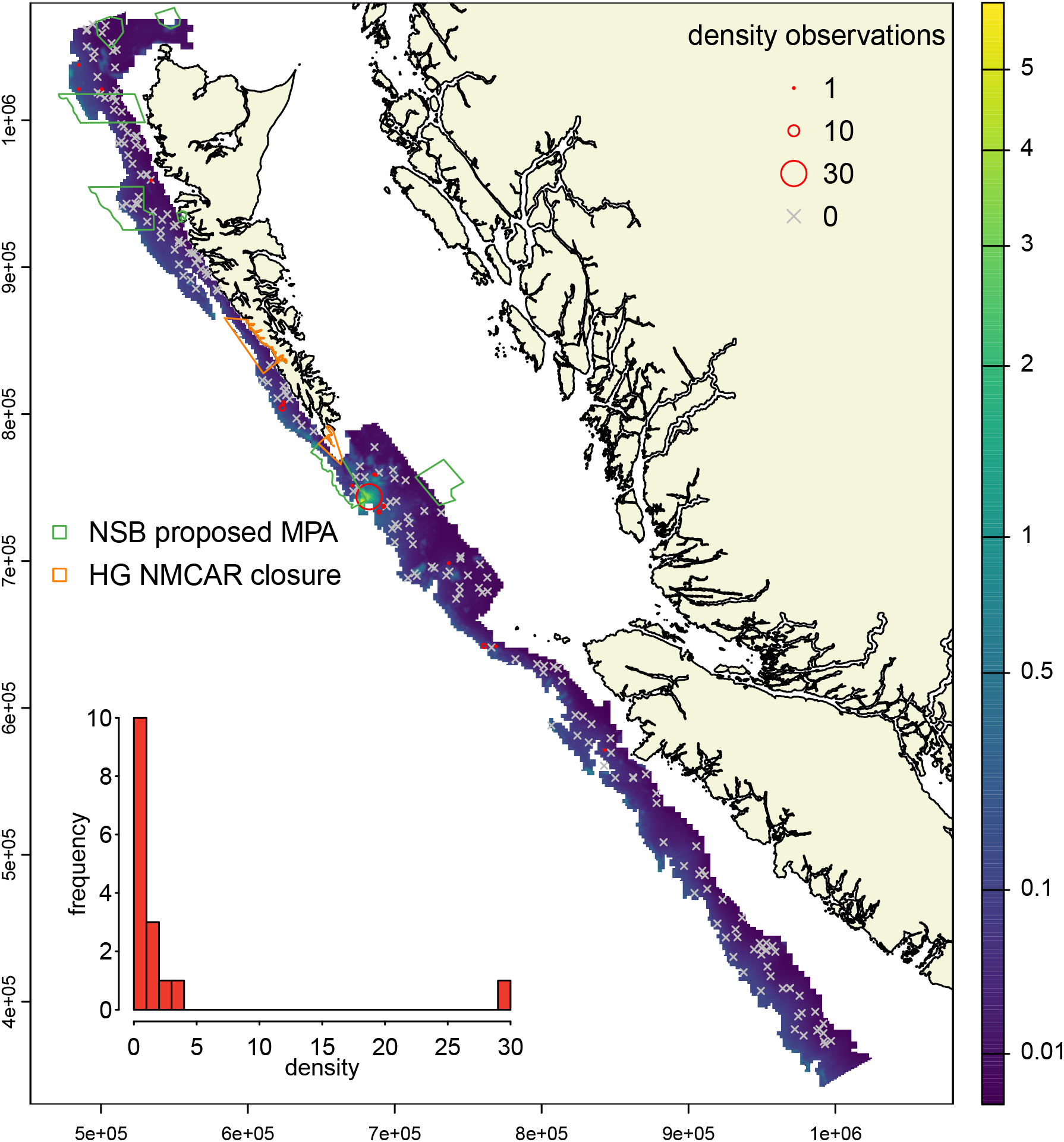
Mean predicted density for corals from spatial models within Sablefish fishing areas in BC. Density (counts per video) for presence observations are shown as red circles sized in proportion to density, while absence observations are shown as grey Xs. We use a cubic root to transform the color scale so that variability in lower density areas can be seen. Coastline data are from Wessel and Smith 1996 and map projection is NAD83 BC Albers (EPSG:3005).

We had 16 presence and 266 absence observations for corals (Fig. 2), including *Heteropolypus* sp., *Isidella* sp., *Paragorgia* sp., *Parastenella* sp., and *Swiftia simplex*. Most observations included relatively low densities with only 1–3 coral colonies, except for one location south of Cape St. James where 30 coral colonies (Alcyonacea and *Parastenella* sp.) were observed on a single camera deployment.

### 3.2. Sablefish fishery footprint

We estimate gear-habitat contact (i.e., footprint) from historical Sablefish fishing within 1353 grid cells of 4 km x 4 km (21 648 km^2^), which corresponds to cells fished by at least 3 unique vessels using longline trap or hook gear from 1990–2024 (Fig S.6). This footprint omits 2 % of the total fishery sets from 1990–2024 that occur in grid cells fished by fewer than three unique vessels that were excluded due to regulations outlined in Canada’s Access to Information and Privacy Act (i.e., the rule of three, Tomasic, 2023). We estimate that 89.9% (95% CI: 86.3–93.8) of these Sablefish fishing grounds have zero gear-habitat contact and that another 8.1% (95% CI: 5.4–10) of those areas experience only one habitat contact event from 1990 to 2024 (Table 3). Estimates of habitat area contacted more than once by fishing gear account for only 2.1 % (95% CI: 0.8–3.7) of fishing grounds.

When considering gear contact in each 4 km x 4 km grid cell, we found that less than 10% of habitat within a grid cell is contacted by fishing gear for the majority (70%) of grid cells (Table S.1). In contrast, we found 1.9% of grid cells are expected to have 50–92% of their habitat contacted by fishing gear from 1990-2024 (Table S.1). The estimates of historical trap and hook effort from 1965–1989, prior to reliable logbook information and gear deployments, are provided in supplementary materials (Fig. S.7–S.8)

#### 3.3. Species Distribution Models

Presence-absence (Figs S.2–S.3) and density (Figs 1–2) models show similar regions with high probability and density for sponges and for corals, respectively. Areas with high probability and density of sponge habitats are concentrated in patchy distributions in northwest, west, and southern Haida Gwaii, at the western end of the Moresby trough within Queen Charlotte Sound, and northwest Vancouver Island (Figs. 1, 3, and S.2). In comparison there are far fewer areas with high probability and density of corals, with the largest habitat areas occurring south of Haida Gwaii (Cape St. James) and to the northwest of Haida Gwaii in deeper waters (Figs. 2, 4, and S.3).

**Figure 3:**
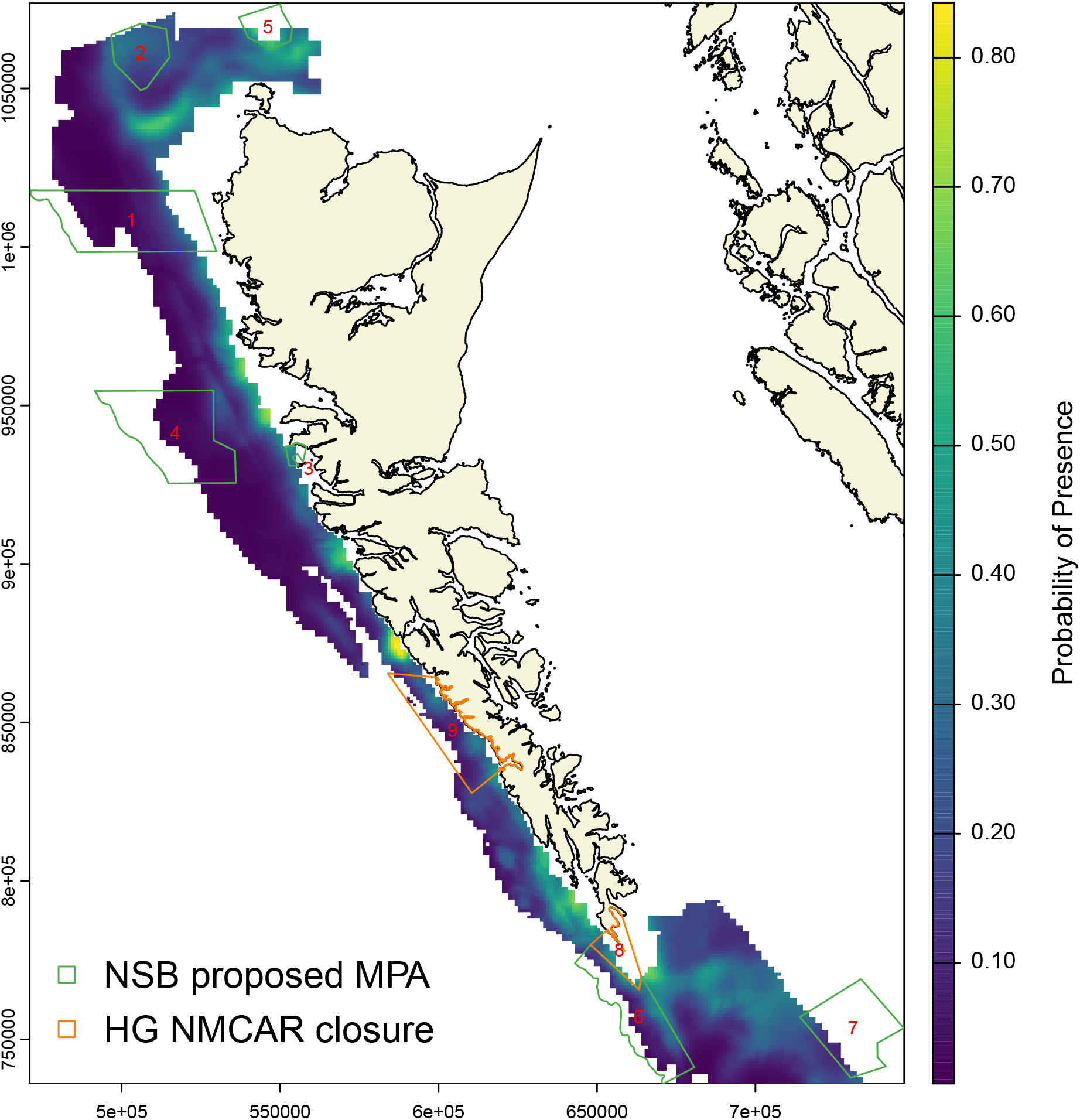
Mean predicted probability of presence for sponges (Porifera) within Sablefish footprint and MPAs. Coastline data are from Wessel and Smith 1996 and map projection is NAD83 BC Albers (EPSG:3005). MPAs are numbered as follows: 1. NSB Zone 502, 2. NSB Zone 506, 3. NSB Zone 483, 4. NSB Zone 503, 5. NSB Zone 501, 6. NSB Zone 505, 7. NSB Zone 510, 8. GH NMCAR South Kunghit Island, 9. GH NMCAR Kwoon Cove to Gowgaia Bay.

**Figure 4:**
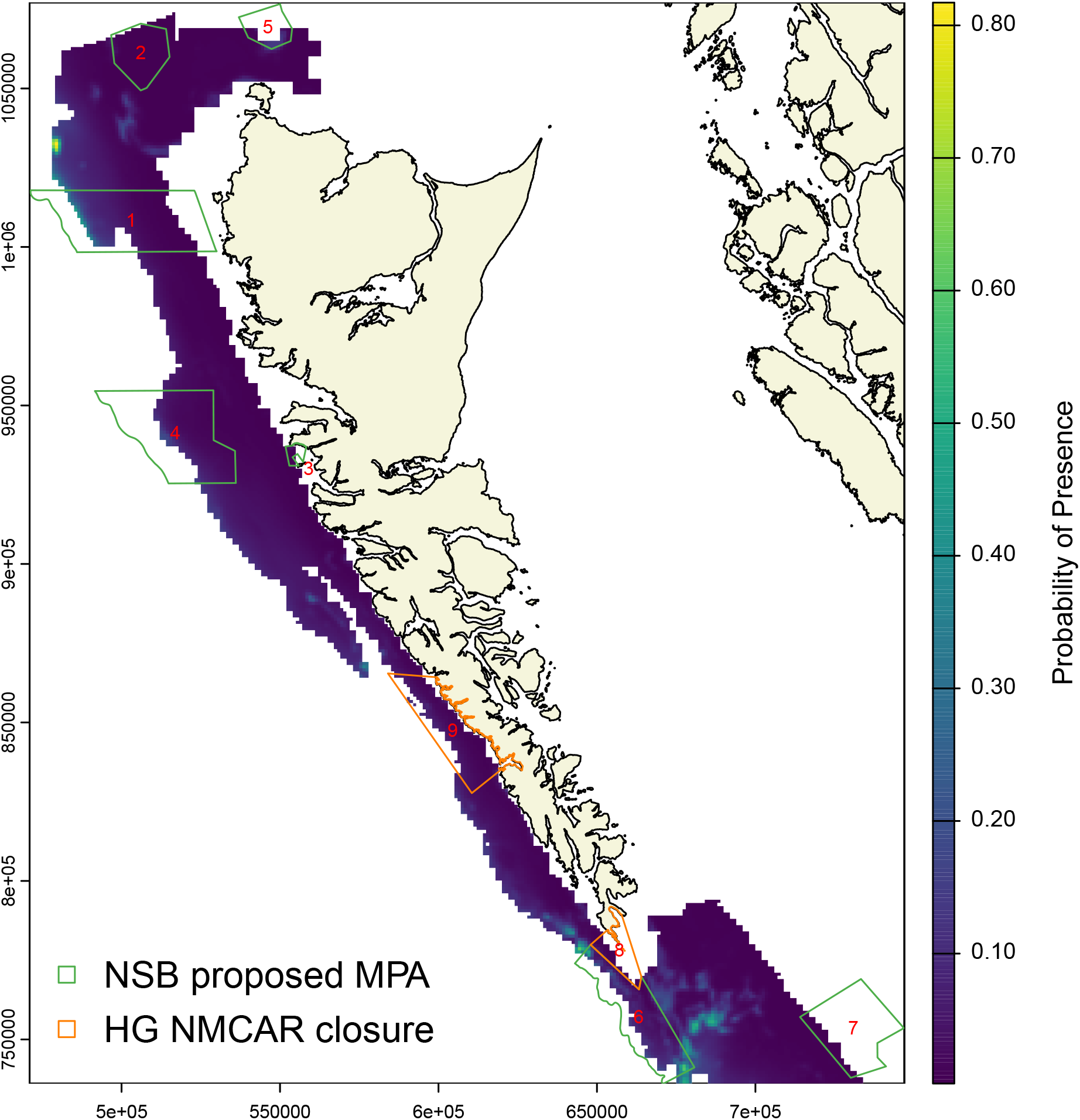
Mean predicted probability of presence for corals (Alcyonacea) within Sablefish footprint and MPAs. Coastline data are from Wessel and Smith 1996 and map projection is NAD83 BC Albers (EPSG:3005). MPAs are numbered as follows: 1. NSB Zone 502, 2. NSB Zone 506, 3. NSB Zone 483, 4. NSB Zone 503, 5. NSB Zone 501, 6. NSB Zone 505, 7. NSB Zone 510, 8. GH NMCAR South Kunghit Island, 9. GH NMCAR Kwoon Cove to Gowgaia Bay.

Performance metrics from 5-fold cross validation presence-absence models (Figs. S.2–S.3) produced mean AUCs of 0.68 (mean SD: 0.12) and 0.77 (mean SD:0.10) for sponges and corals, respectively, indicating acceptable performance (DeLong et al., 1988). Covariate relationships for probability of presence or both coral and sponge models are provided in supplementary material (Figs S4-S5).

### 3.4. Coral and sponge habitat overlap with fishery closures

We identified three MPAs closed to Sablefish fishing within the GH NMCAR and eight proposed MPAs within the NSB with coral or sponge conservation objectives that overlapped with the modelled habitat area (Figs. 3–4). We excluded two sites (Queen Charlotte Sound Zone 511 in NSB, Sgaan Gwaay in GH NMCAR) from further analysis because both have less than 4% (2–4 km^2^) of their area within the boundaries of the Sablefish footprint.

The proportion of predicted habitat areas within each MPA (Table 2) ranged from 3.1–46.9% (5.5–63.4 km^2^) for sponges and 0.4–9.8% (0.2–35.8 km^2^) for corals (Figs. 3–4). We estimated that 15.6% and 3.1% of areas open to Sablefish fishing contained sponge and coral habitats, respectively. We found that one MPA in GH NMCAR (GH NMCAR South Kunghit Island MPA) and three proposed MPAs in NSB (NSB Zones 483, 501, and 506) contained a higher proportion of sponge habitat (24.4-46.9%) relative to that in open areas, whereas two NSB Zones (502, 503) had much less (3-8%). Three areas (GH NMCAR South Kunghit Island MPA, NSB Zone 483, NSB Zone 501) had similar proportions of sponge habitat (16.7-18.3%) relative to open areas. For corals, we found higher proportions of coral habitat (6.0-9.8%) in three proposed MPAs (NSB Zones 501, 502, 505) relative to open areas, while three NSB zones (483, 506, 510) had less (0.4-1.1%). There were another three areas (GH NMCAR South Kunghit Island, GH NMCAR Kwoon Cove to Gowgaia Bay, NSB Zone 503) with similar proportions of coral habitat (1.6-3.8%) relative to open areas.

**Table 2:** Predicted sponge and coral habitat within the Sablefish fishery footprint for locations inside and outside MPAs (current and proposed). The total overlap area between MPAs and the footprint are shown in the 2nd column, along with the amount of coral and sponge habitat in each location as area and percentage. The final row shows the total area and habitats inside the footprint that are open to fishing (i.e., excluding MPAs in rows 1-9).

| Location | Area/Habitat in Fishing Grounds |  |  |  |  |
| --- | --- | --- | --- | --- | --- |
|  | Total (km<sup>2</sup>) | Sponges (km<sup>2</sup>) | Corals(km<sup>2</sup>) | % Sponge | % Coral |
| NSB Zone 502 | 559.9 | 44.1 | 33.6 | 7.9 | 6.0 |
| NSB Zone 506 | 259.9 | 63.4 | 1.2 | 24.4 | 0.4 |
| NSB Zone 483 | 15.1 | 5.5 | 0.2 | 36.3 | 1.1 |
| NSB Zone 503 | 470.5 | 14.7 | 18.1 | 3.1 | 3.8 |
| NSB Zone 501 | 43.3 | 20.3 | 2.6 | 46.9 | 6.0 |
| NSB Zone 505 | 364.8 | 62.7 | 35.8 | 17.2 | 9.8 |
| NSB Zone 510 | 71.1 | 11.8 | 0.3 | 16.7 | 0.4 |
| GH NMCAR South Kunghit Island | 44.4 | 16.6 | 0.7 | 37.3 | 1.6 |
| GH NMCAR Kwoon Cove to Gowgaia Bay | 340.0 | 62.1 | 10.0 | 18.3 | 2.9 |
| Open fishing area | 21412.9 | 3337.0 | 657.6 | 15.6 | 3.1 |

**Table 3:**
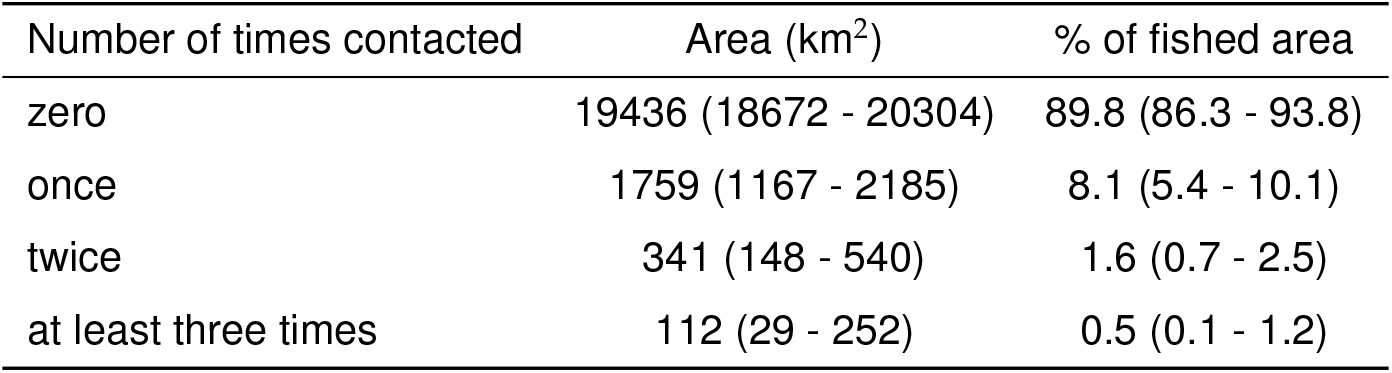
Frequency of habitat contact by Sablefish longline trap and hook fishing gear from 1990-2024 within the total Sablefish fishing grounds of 21 648 km^2^ (1353 grid cells of 4 km x 4 km). Mean estimates are shown with 95% CIs in brackets.

| Number of times contacted | Area (km<sup>2</sup>) | % of fished area |
| --- | --- | --- |
| zero | 19436 (18672 - 20304) | 89.8 (86.3 - 93.8) |
| once | 1759 (1167 - 2185) | 8.1 (5.4 - 10.1) |
| twice | 341 (148 - 540) | 1.6 (0.7 - 2.5) |
| at least three times | 112 (29 - 252) | 0.5 (0.1 - 1.2) |

### 3.5. Relative benthic habitat status

We estimate that the aggregated RBS for sponge habitats within BC Sablefish fishing grounds decline from assumed unfished levels with RBS = 1 to a mean RBS of 0.96 (95% CI: 0.93–98) in 2024, accounting for habitat contact from Sablefish longline trap and hook gears (Fig. 5). The largest habitat status declines occured between 1967 and 1979 in the early fishing years with the highest longline catch and estimated historical fishing effort (Figs S.7–S.8), during which RBS decreased to 0.93 (95% CI: 92–95).

**Figure 5:**
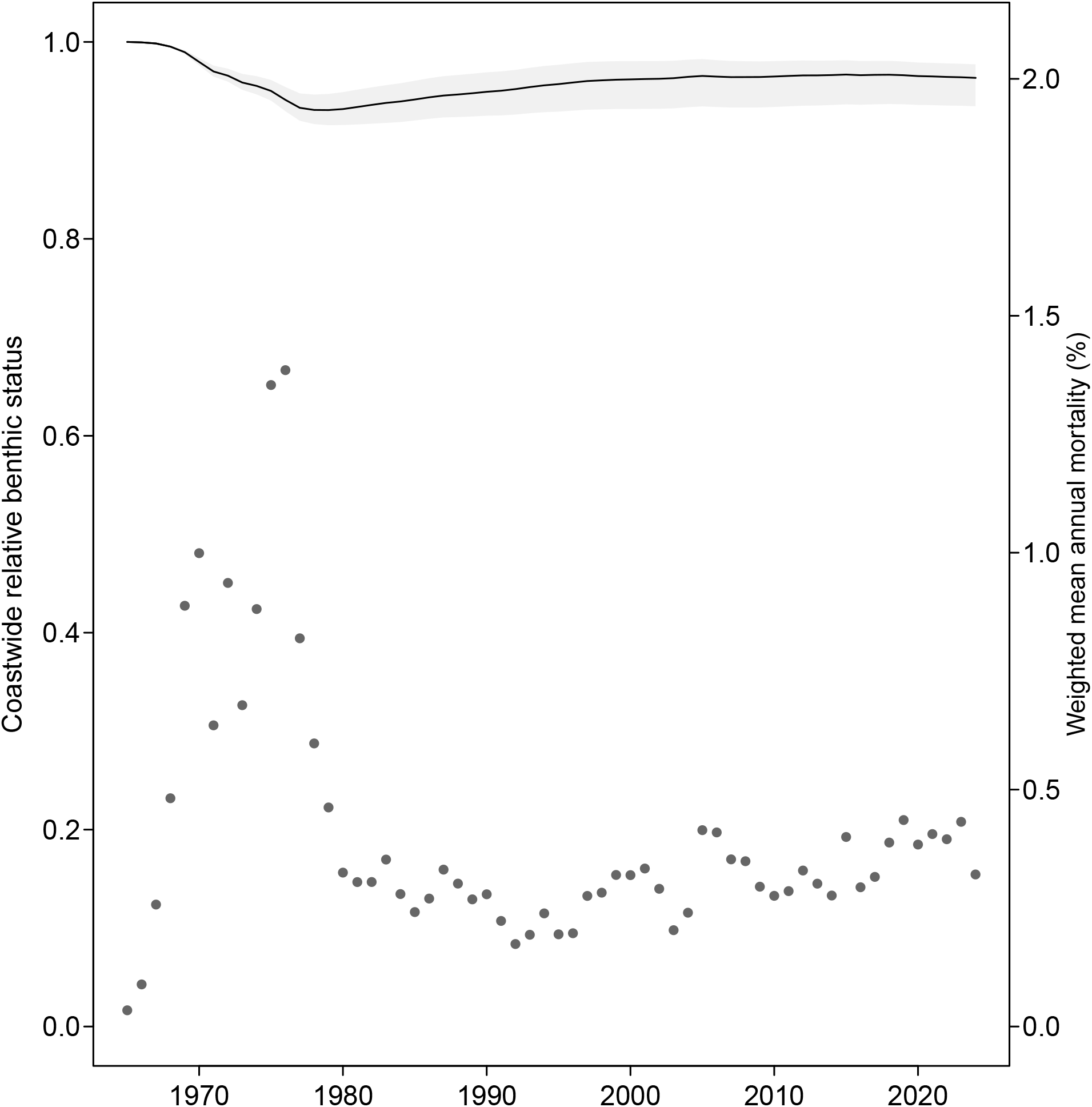
Relative benthic status (RBS) for aggregated sponge habitats within BC Sablefish fishing grounds from 1965-2024, accounting for habitat contact from Sablefish longline trap and hook gears. The solid black line is the mean RBS, while grey polygons indicate 95% confidence intervals. Grey dots are mean annual mortality 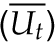 from Sablefish longline trap and hook gear, weighted by biomass in each grid cell, shown on the second y-axis on right side.

We found that mean RBS in 2024 varied from 0.71 (95% CI:0.18–0.98) to 1 (95% CI:0.999–1) across all *I* = 25 081 1 km x 1 km grid cells examined (Fig. 6, S.9–S.10), variation which is primarily driven by spatial differences in fisheries effort. We found that 99% of grid cells (24 913 km^2^) have mean RBS greater than 0.90, while only 0.1% (16 km^2^) have RBS between 0.71 and 0.80 (Table 4).

**Table 4:** Summary of mean RBS for sponge habitats in 2024 across 25 081 grid cells (1 km x 1 km) considered in risk assessment. Higher RBS values indicate less decline in habitat status from 1990 to 2024.

|  | Mean RBS | n cells | Area (km<sup>2</sup>) | % of cells |
| --- | --- | --- | --- | --- |
| 8 | 0.71 - 0.8 | 16 | 16 | 0.1 |
| 9 | 0.8 - 0.9 | 152 | 152 | 0.6 |
| 10 | 0.9 - 1 | 24913 | 24913 | 99.3 |

**Figure 6:**
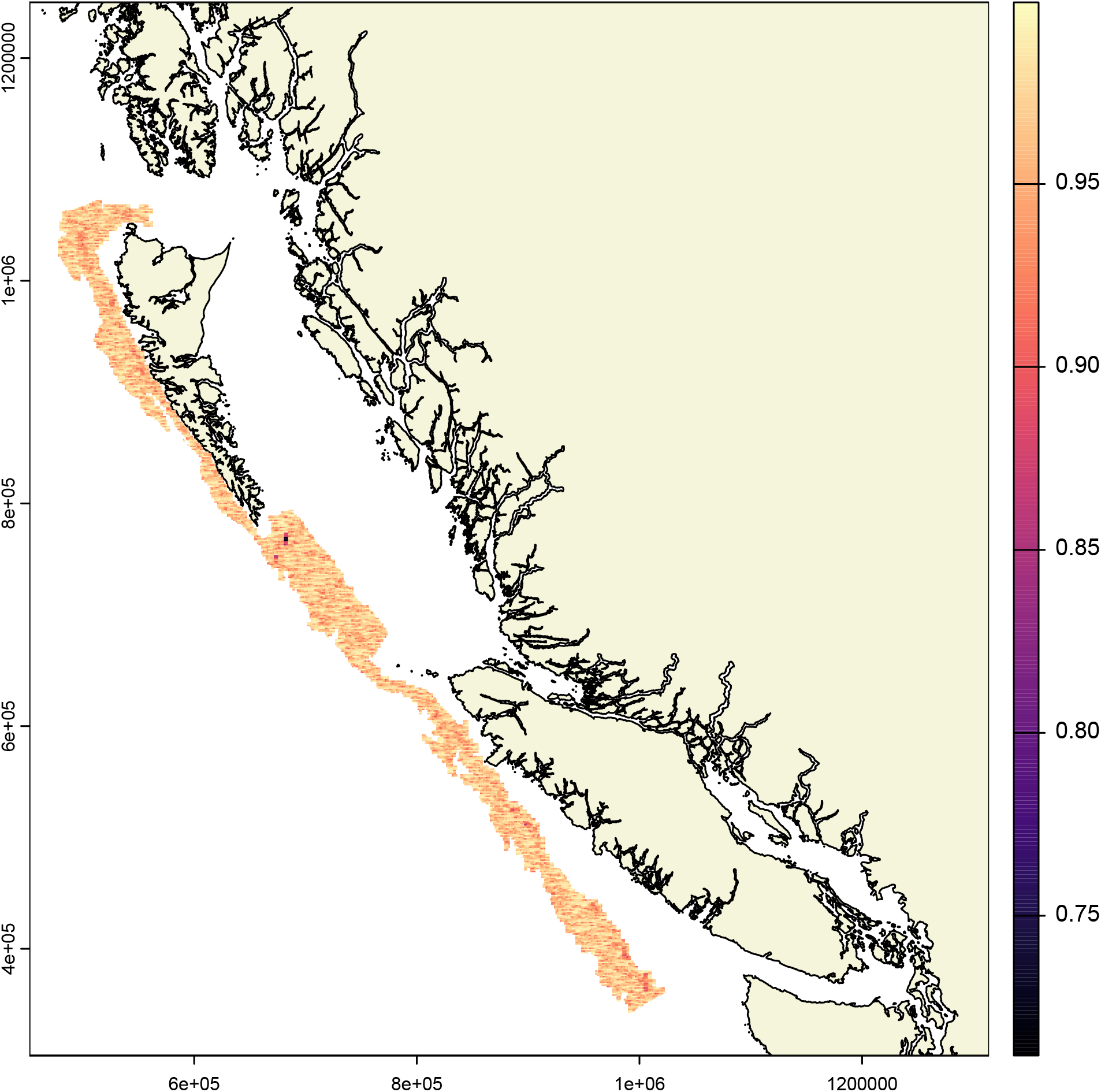
Mean RBS for sponge habitats in 2024 for *i* = 25 081 grid cells (1 km x 1 km) examined. Coastline data are from Wessel and Smith 1996 and map projection is NAD83 BC Albers (EPSG:3005).

## 4. Discussion

We demonstrated a quantitative approach for assessing risks from bottom longline trap and hook fisheries on seafloor habitats, focusing on sponge habitats and the BC Sablefish fishery as a case study. We used relative benthic status as a measure of risk, estimating that mean RBS for total sponge habitats in BC Sablefish fishing grounds declined from 1 to 0.96 due to Sablefish fishing from 1965 to 2024. This decline is a 4% reduction in habitat status compared to pre-fishery levels. The majority of the habitat decline occurred between 1967 and 1979 during the period of peak catch and effort from the longline hook fishery(Figs S.7–S.8), when RBS values indicated a 7% reduction in habitat status. 0ur risk assessment indicates some habitat recovery from 1980 to 2024 (increasing RBS), coinciding with reduced fishing effort. The 2024 habitat status at finer spatial scales of 1 km^2^ grid cells varies depending on the concentration of fishing effort. The majority of grid cells present similar trends to the coastwide aggregate status with habitat declines less than 10% (RBS > 0.9) since 1965 for 99% of grid cells examined (Table 4). The most impacted grid cells had slightly larger habitat declines of 20 to 29% (0.71 <RBS < 0.8), occurring in a small proportion of the fishing grounds (0.1% of grid cells) South of Haida Gwaii (Fig. S.11). Our analysis provides fine-scale information on habitat impacts from fishing gear, providing key information for conservation planning and fisheries management.

Our quantitative risk assessment approach (Doherty et al., 2025) involves five key steps: i) data collection, ii) quantifying habitat contact from fishing gear, iii) mapping SBHs, iv) estimating habitat status relative to pre-fishery levels via fine-scale spatial logistic population models, and v) evaluating economic and conservation trade-offs under alternative management measures designed to protect SBHs. The model-based sponge density estimates from (iii) are used to initialize population models in (iv) in 1km^2^ grid cells, while accounting for annual mortality from fishing gear contact from (ii). Species distribution models for coral and sponge habitats used for step ii) reveal key areas of overlap between Sablefish fishing grounds and spatial fishery closures designed to protect habitat. Notable areas of fishing ground overlap with sponge habitats occur to the south of Cape St James, northwest Haida Gwaii, and northwest Vancouver Island. In contrast, coral habitats were rarely observed in the Sablefish fishing grounds, occurring in only 6% of survey sets, and when observed were typically low densities of 1-2 colonies. There was one location where 30 coral colonies were observed to the south of Cape St. James, which occurs near high densities of historical Sablefish fishing effort (Figs. 2, S.6). These maps of coral and sponge habitat density can inform candidate management measures that allow trade-offs in conservation and economic performance to be evaluated when considering marine closures in BC (step v).

The results of our risk assessment (step iv) can inform management strategies designed to protect habitats, providing useful information for evaluating existing and proposed MPAs in BC. Results from our habitat models and risk assessment could be used to identify habitat areas of higher risk from fishing gear impacts to inform spatial closure areas that reduce overall risk to coastwide or localized habitats. Notably, our habitat models indicate several MPAs contain large areas with low probability of coral and sponge habitats. In particular, we found low probability of sponge and coral habitats for large areas of currently proposed NSB closures (NSB Zones 502, 503, and 505, Figs. 3 - 4), which is ground-truthed by multiple absence observations from trap-cameras (Figs. 1 - 2). Similarly, results from our risk assessment highlight large areas of low habitat risk within proposed MPAs, as well as areas of higher risk outside of proposed MPAs. For example, the 16 km^2^ with the greatest habitat impacts (RBS of 0.73 to 0.8) occur south of Cape St James (Figs. 6) in an area that overlaps with high densities of observed and predicted coral and sponge habitats (Figs. 1 - 2). These grid cells occur outside of currently proposed MPAs, to the east of NSB Zone 505 (Fig S.11). Similarly, grid cells in the other current and proposed MPAs include habitats identified as low risk (RBS > 0.85). Ideally future decisions for MPAs designed to protect seafloor habitats will include habitat models and spatial risk assessment, such as those provided in this paper, and allow for adjustments to boundaries over time to reflect new information on habitat distribution and fishing risks.

Step (v) of our risk assessment involves evaluating expectations for future habitat status and fishing effort, and considering trade-offs from alternative management choices. Our risk assessment approach identifies fishing grounds at the 1 km^2^ resolution that have the greatest and lowest habitat impacts, which can be used to evaluate trade-offs between conservation and economic outcomes associated with spatial closures and displaced fishing effort. Low risk areas within MPAs could be considered for reopening to fisheries that are capable of quantifying and monitoring their fishing impacts. For example, three of the currently proposed NSB closures (NSB Zones 502, 503, and 510, Table 2) contained a smaller proportion of sponge habitats than the proportion predicted for areas open to fishing. Such closures could be counter-productive to conservation goals, by shifting fishing effort and increasing fishing impacts into remaining open fishing areas with more sponge habitats. If closures do not contain higher densities of SBHs than densities in fishing areas, they have potential to reduce aggregate coastwide RBS even while improving RBS within closed areas. Furthermore, increased fishery closures reduce available fishing grounds, which can concentrate fishing effort in smaller areas, creating localized depletion and leading to reduced catch rates for target species. In this case, more fishing gear must be deployed in a smaller area to achieve the same catch relative to the status quo prior to any closures. This scenario highlights the spatial trade-offs associated with managing seafloor habitats that are not well understood or studied, whereby conservation goals may need to clearly articulate the spatial scale for habitat objectives and whether they aim to prioritize habitat status in protected areas, even if it means greater impacts outside of closures. Our habitat risk assessment can be integrated into management strategy evaluation (MSE, Punt et al., 2016) to evaluate the effectiveness of alternative habitat conservation strategies for achieving habitat conservation goals. Examples of such strategies can include testing alternative boundaries for fisheries closures or MPAs, move-on rules, and coral and sponge bycatch limits (Parker et al., 2009; Penney et al., 2009; Wallace et al., 2015a).

There is potential to improve maps of habitat distribution for management planning and future risk assessment of seafloor ecosystems in BC. In this paper, we developed habitat distribution models for presence-absence and density of coral and sponges, using deep-water camera observation collected during the BC Sablefish Stratified Random Survey from 2013-2017. While these observations provide a reasonable number of data points for fitting presence-absence models (n=282), the low prevalence of corals (6%) and sponges (10%) among camera observations meant we had low sample sizes for fitting density models that use only the presence observations. Future work could consider new sampling designs or data types to improve density estimates for coral and sponge species and mapping areas with high density. This might include techniques such as adaptive cluster sampling that target areas with higher probabilities of presence to improve accuracy in density estimates (Thompson, 1990; Bowering et al., 2018; Piazza et al., 2020). Similarly, additional sampling could be directed to areas where density predictions from SDMs have high uncertainty, possibly due to data limitations. The number of presence observations could also be increased by fitting a combined sponge and coral habitat model, since observed coral and sponge locations commonly overlapped. Additionally, future habitat modelling could combine our presence-absence data with other coral and sponge data for BC (Nephin et al., 2023), including opportunistic presence-only data from fisheries bycatch records, corrected for biased sampling (Fithian et al., 2015). For example, estimates of catch per unit effort for coral and sponges can provide reliable abundance indices for habitat models (Rooper et al., 2011). Our density observations are in counts per video since we do not have reliable estimates of the camera field of view from each camera observation. Future camera systems for habitat sampling could incorporate new sensors or technologies (e.g., lasers, stereo cameras) that allow estimating field of view (Williams et al., 2014; Rooper et al., 2016), enabling improved estimates of density (e.g. counts per km^2^) that can be directly converted to biomass by multiplying density by area.

Future iterations of habitat risk assessment in BC could be improved in at least four ways. One of the key uncertainties for coral and sponge risk assessment is their recovery rates following damage, for which there are few estimates available. Here we use estimates for sponges based on models from trawl surveys in Alaska that predict it would take 20 years (95%CI: 13-36) to recover to 80% of initial biomass levels following a 67% mortality event (Rooper et al., 2011). Such recovery rates are likely taxa specific and therefore future analyses could consider the composition of sponge taxa in a given area to inform choices for recovery rates; however, there remain few studies on deep-water coral and sponge population dynamics. In the absence of improved data for sponge and coral recovery rates, alternative scenarios or additional sensitivity analyses for intrinsic growth rates could be considered. Second, our risk assessment only accounts for Sablefish fishing impacts on habitats and does not consider other BC fisheries. A more robust risk assessment would include cumulative fishing impacts from all bottom-contact fisheries that overlap with Sablefish fishing grounds, such as other longline (Pacific Halibut, Rockfish), trap (Prawn), and groundfish bottom trawl fisheries (Wallace et al., 2015b). Third, there remains considerable uncertainty about historical numbers of traps and hooks deployed by the Sablefish fishery prior to 1990, for which we developed a reconstructed effort time series. The longline hook effort in the 1960s and 1970s is responsible for the majority of the predicted sponge habitat loss in our risk assessment. Alternative reconstructed effort time series for the 1965-1989 period could be explored as part of future risk assessments for the Sablefish fishery. Finally, there is a lack of quantitative objectives for coral and sponge habitats in Canada, which makes it challenging to evaluate if aggregate and spatial RBS scores from our risk assessment are achieving conservation goals. Habitat objectives should be “specific, measurable, achievable, realistic and time-bounded (SMART) objectives” and “identified at smaller spatial scales” (of Canada, 2014). It is widely recognized that clear quantitative objectives are needed for achieving ecosystem and biodiversity goals (Hilborn et al., 2004; Tear et al., 2005; Agardy et al., 2016), without which it is difficult to evaluate the performance of alternative management strategies with respect to tradeoffs between conservation and economic benefits that provide food production and employment.

## 5. Conclusion

We have demonstrated a quantitative risk assessment framework to evaluate the impacts of longline fishing gear on seafloor habitats, providing metrics for evaluating habitat status relative to ecosystem objectives. Our approach uses a standard risk metric (RBS) for evaluating fishing impacts on seafloor habitats that can be broadly applied to bottom fisheries, supporting evidence-based decision making and reducing reliance on qualitative approaches for evaluating fishing risks to VMEs. Such habitat metrics can be incorporated into fisheries management strategy evaluation, allowing resource managers to compare performance of alternative strategies against a broader suite of sustainability objectives that include habitat, fish stocks, and fisheries catch.

## 6. Acknowledgements

We thank the Canadian Sablefish Association and Wild Canadian Sablefish Ltd. for their financial and in-kind support at all stages of this project from the initial design to field deployments and analysis. We especially thank the fishing masters who expertly deployed-retrieved camera traps on commercial fishing sets. This project was also made possible by Kristina Castle and Malcolm Wyeth, who assisted with camera design, data collection and preparation, equipment preparation, and training of at-sea observers. We thank Kendra Holt and Sean Anderson for helpful comments that improved the manuscript.

