## supplementary material for "A quantitative risk assessment approach for longline fishing gear impacts on seafloor habitats"

### S1. Supplementary Material

This supplementary includes:

- Summary of gear-habitat contact estimates for Sablefish longline trap and hook deployments from 1990-2024 within Sablefish fishing grounds (Table S1).
- Maps of uncertainty (95% CIs) for predicted density of corals and sponges (Figure S1)
- Probability of presence maps for corals and sponges in BC (Figures S2-S3)
- Covariate relationships for probability of presence (i.e., marginal effects plots) for coral and sponge models (Figures S4-S5)
- Footprint map showing proportion of habitat contact for Sablefish longline trap and hook fishing from 1990 to 2024 (Figure S6)
- Historical reconstructed CPUE and effort time series for longline trap and hook fishing by the BC Sablefish fishery (Figures S7-S8)
- Uncertainty (0.025 and 0.975 quantiles) in spatial estimates of Relative Benthic Score (RBS) for 2024 for $i$ = 25 081 1 km x 1 km grid cells considered in sponge habitat risk assessment (Figures S9-S10)
- Map of mean RBS for 2024 for sponge habitat risk assessment, including overlap with current and proposed MPAs (Figure S11)

Table S.1: Summary of gear-habitat contact estimates for Sablefish longline trap and hook deployments from 1990-2024 within Sablefish fishing grounds (1353 grid cells of 4 km x 4 km). This excludes 2% of the total fishery sets from 1990-2024 that occur in grid locations fished by fewer than three unique vessels due to regulations outlined in Canada's Access to Information and Privacy (i.e., the rule of three, Tomasic 2023).

| % of cell contacted at least once | n cells | Area (km2) | % of cells | cumulative % of cells |
|---|---|---|---|---|
| 0 - 1 | 371 | 5,936 | 27.4 | 27.4 |
| 1 - 2 | 188 | 3,008 | 13.9 | 41.3 |
| 2 - 3 | 107 | 1,712 | 7.9 | 49.2 |
| 3 - 4 | 62 | 992 | 4.6 | 53.8 |
| 4 - 5 | 53 | 848 | 3.9 | 57.7 |
| 5 - 10 | 171 | 2,736 | 12.6 | 70.3 |
| 10 - 20 | 168 | 2,688 | 12.4 | 82.7 |
| 20 - 30 | 90 | 1,440 | 6.7 | 89.4 |
| 30 - 40 | 51 | 816 | 3.8 | 93.2 |
| 40 - 50 | 34 | 544 | 2.5 | 95.7 |
| 50 - 60 | 33 | 528 | 2.4 | 98.1 |
| 60 - 70 | 13 | 208 | 1.0 | 99.1 |
| 70 - 80 | 9 | 144 | 0.7 | 99.8 |
| 80 - 90 | 0 | 0 | 0.0 | 99.8 |
| 90 - 92 | 3 | 48 | 0.2 | 100.0 |

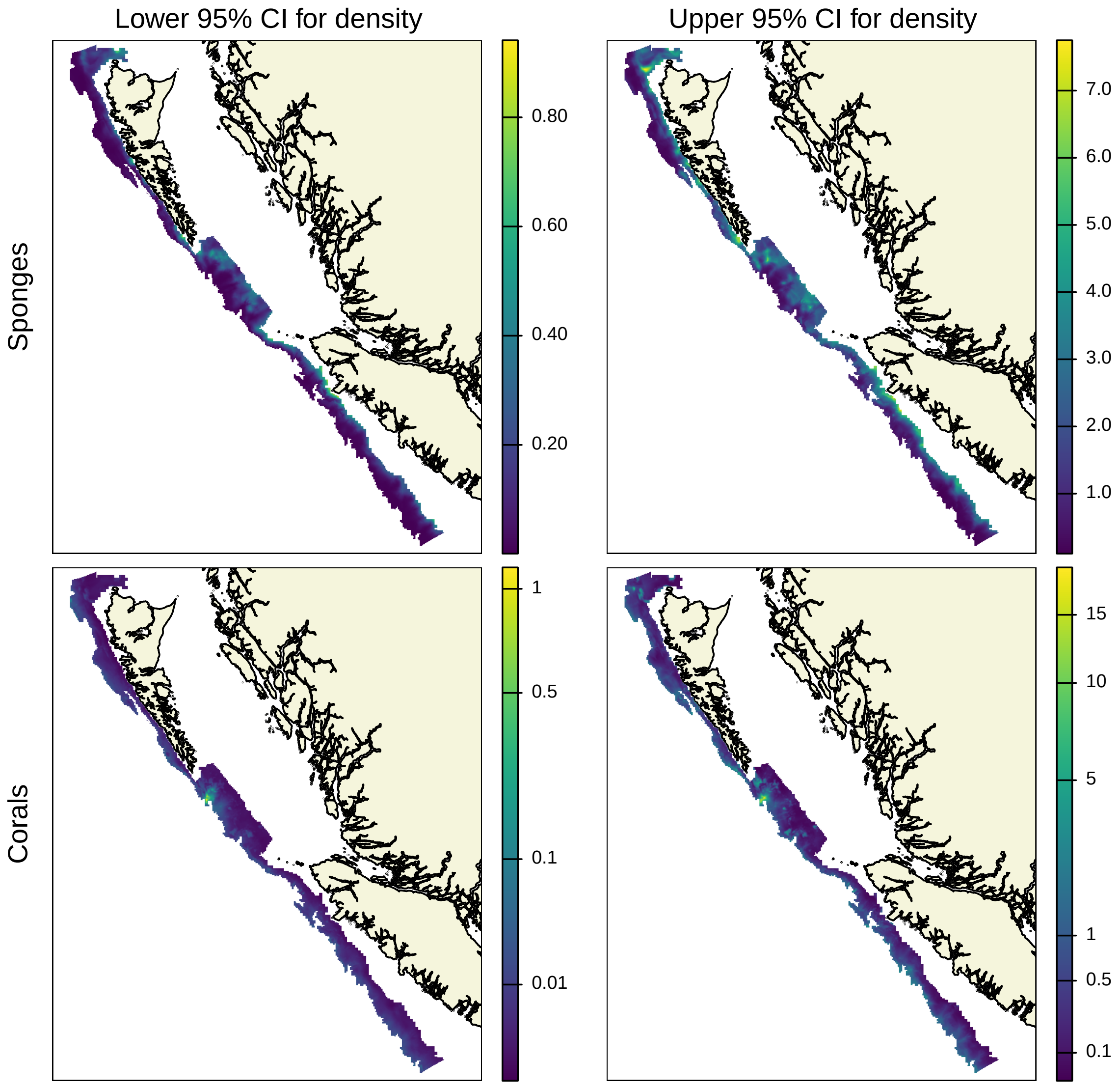


Figure S.1: Lower (left) and upper 95% CIs (right) for predicted density (structures/video) of sponges (top) and corals (bottom) from spatial models within Sablefish fishing areas in BC. We use a cubic root to transform the color scale for corals so that variability in lower density areas can be seen. Coastline data are from Wessel and Smith 1996 and map projection is NAD83 BC Albers (EPSG:3005).

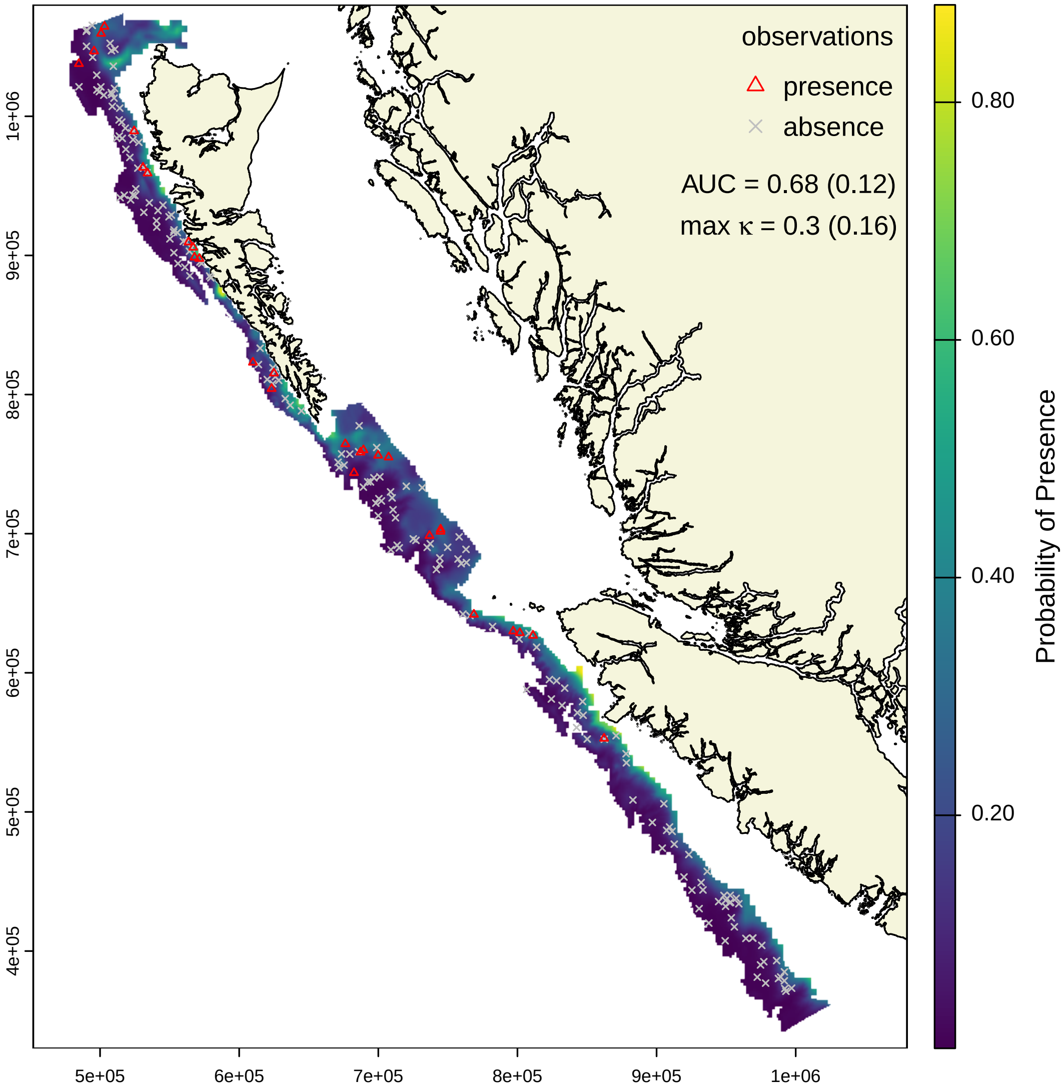


Figure S.2: Mean predicted probability of presence for sponges (Porifera) from spatial models within Sablefish fishing areas in BC. Coastline data are from Wessel and Smith 1996 and map projection is NAD83 BC Albers (EPSG:3005).

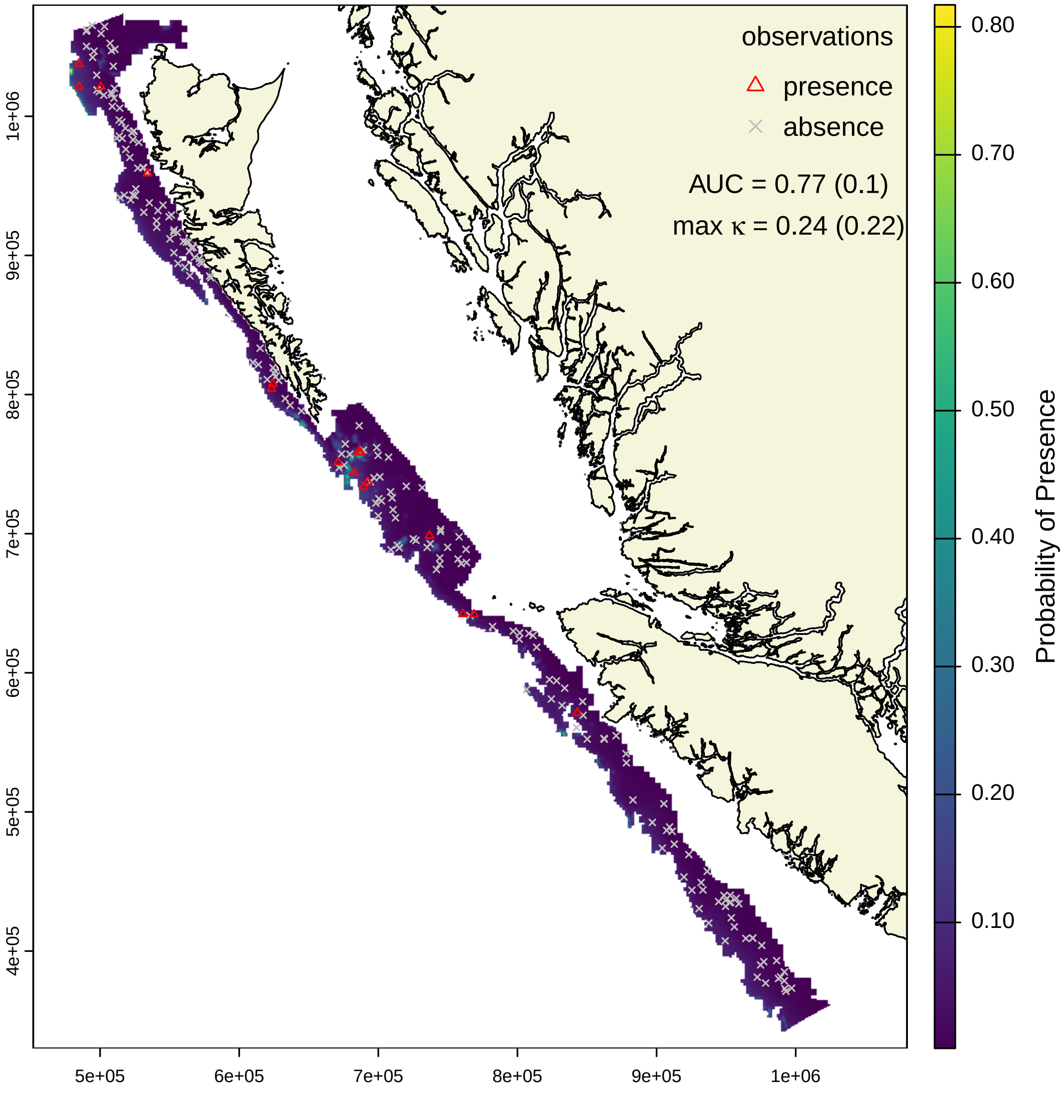


Figure S.3: Mean predicted probability of presence for corals (Alcyonacea) from spatial models within Sablefish fishing areas in BC. Coastline data are from Wessel and Smith 1996 and map projection is NAD83 BC Albers (EPSG:3005).

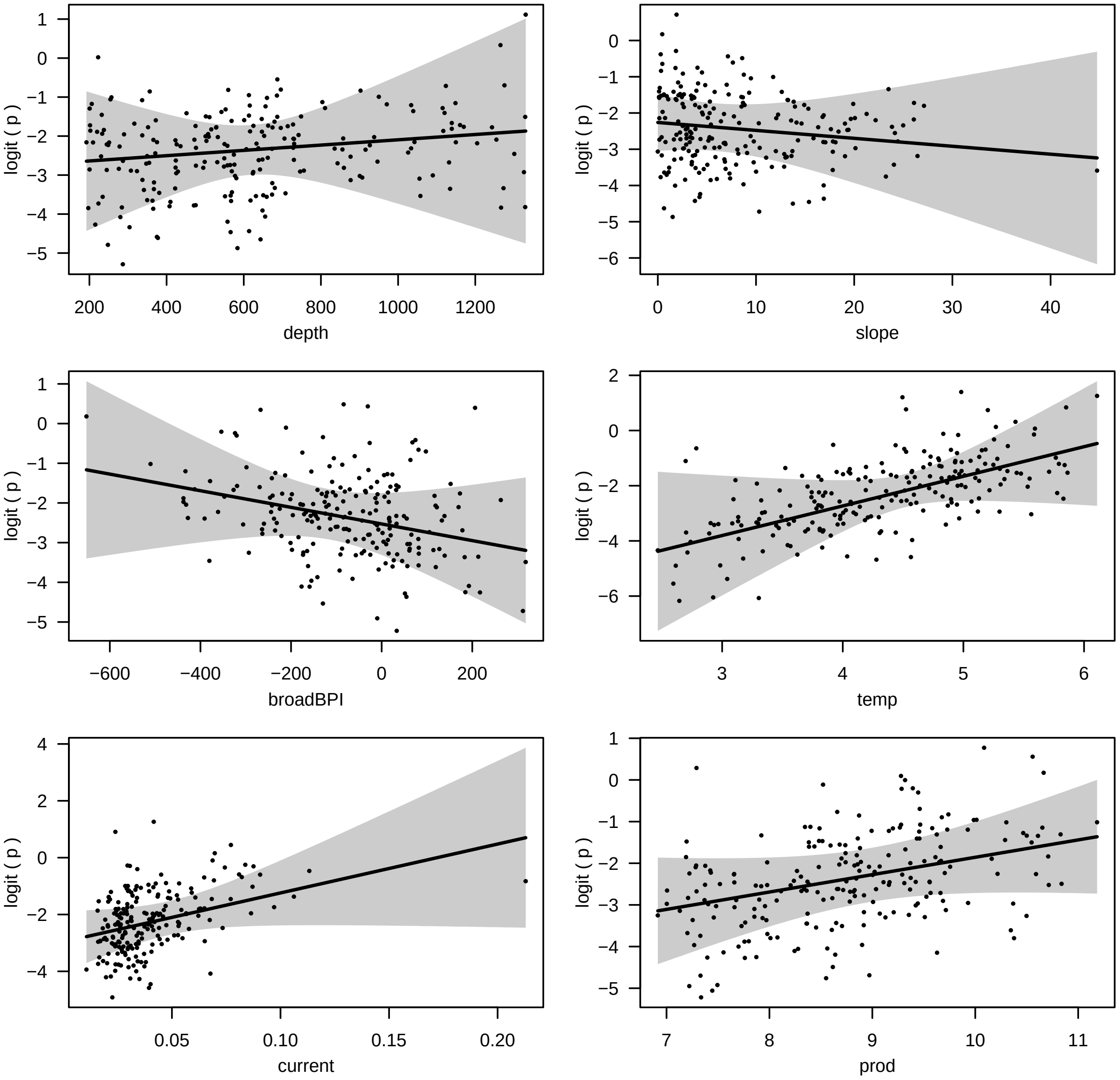


Figure S.4: Conditional effects of covariates on logit probability of sponge presence, including mean predictions (black line), 95% CIs (grey polygon), and partial residuals (black dots).

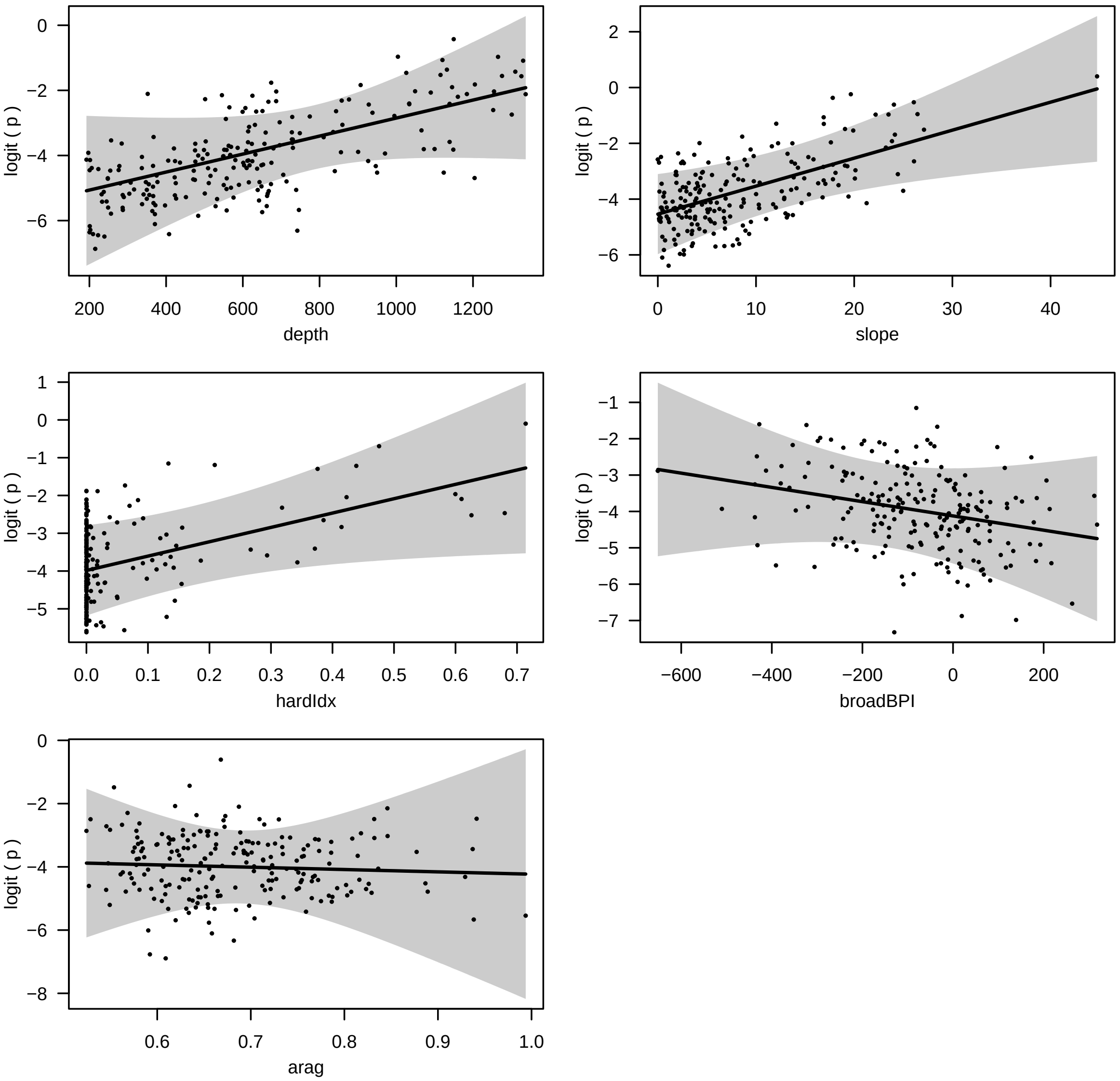


Figure S.5: Conditional effects of covariates on logit probability of coral presence, including mean predictions (black line), 95% CIs (grey polygon), and partial residuals (black dots).

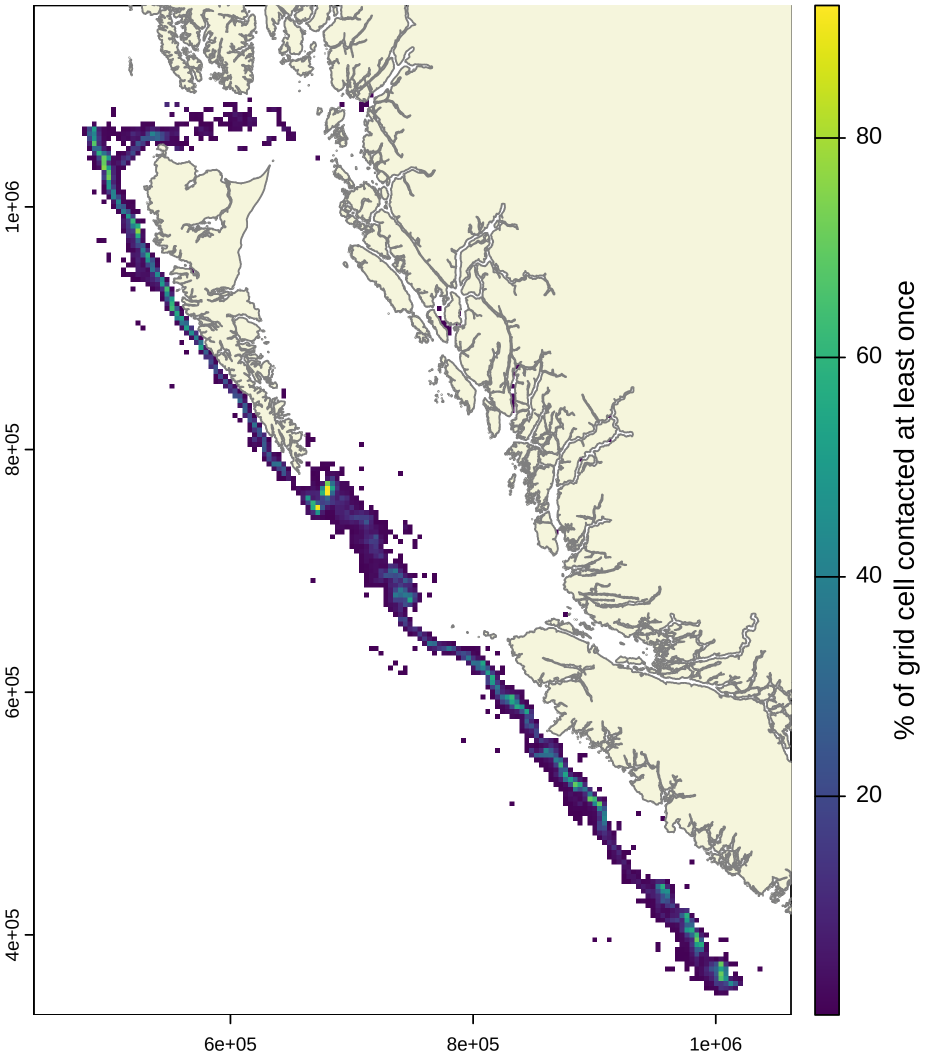


Figure S.6: Proportion of 4 km × 4 km grid cells contacted by Sablefish longline trap or hook gear at least once from 1990 to 2024. Note that this excludes 2% of the total fishery sets, which occur in grid locations fished by fewer than three vessels. Coastline data are from Wessel and Smith 1996 and map projection is NAD83 BC Albers (EPSG:3005).

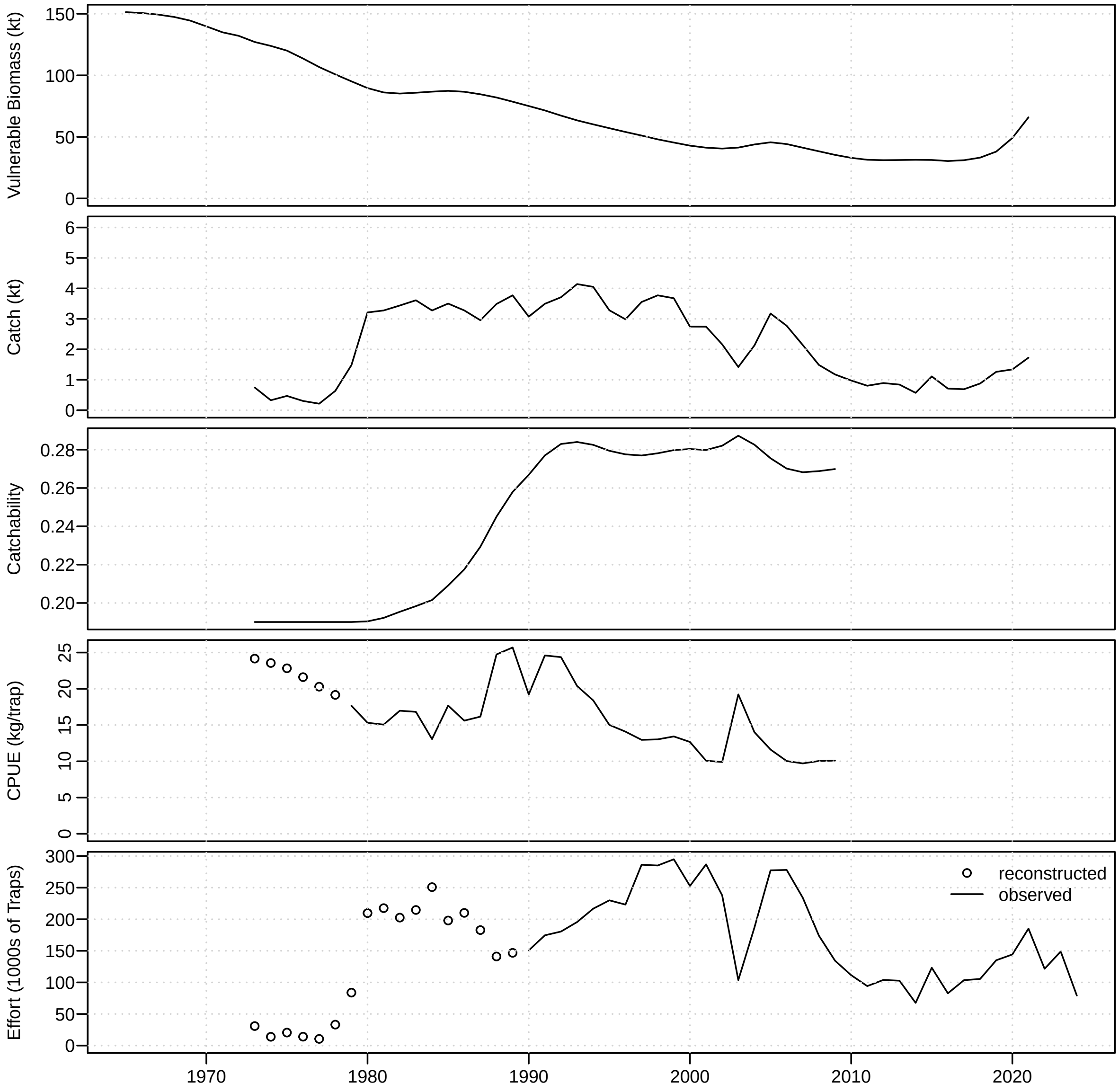


Figure S.7: Vulnerable biomass (1st panel), catch (2nd panel), catchability (3rd panel), CPUE (4th panel), and effort (5th panel) from the BC Sablefish commercial longline trap fishery from 1965–2024. Reconstructed (open circles) and observed (solid line) values are shown in the bottom panels for CPUE and effort. Biomass (1965–2021) and catchability (1973–2009) are estimates from the BC Sablefish operating model.

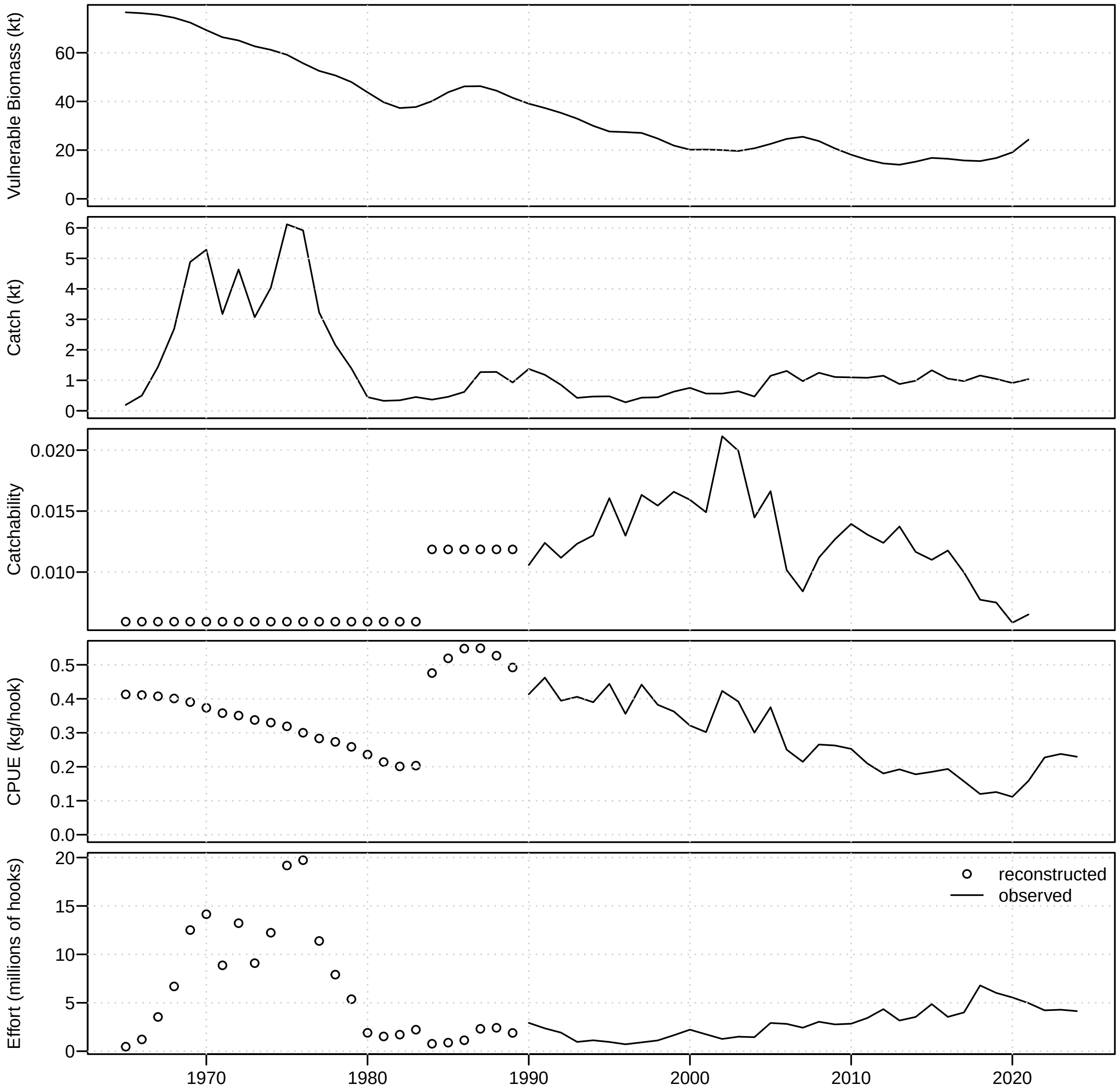


Figure S.8: Vulnerable biomass (1st panel), catch (2nd panel), catchability (3rd panel), CPUE (4th panel), and effort (5th panel) from the BC Sablefish commercial longline hook fishery, including directed Sablefish and Sablefish/Halibut combined trips, from 1965–2024. Reconstructed (open circles) and observed (solid line) are shown in bottom panels for CPUE and effort. Catchability from 1990 to 2021 (solid line) is estimated from biomass and CPUE time series, while fixed values are used for 1965–1983 and 1984–1989 that reflect historical fishing conditions (see Step 2 in methods). Biomass (1965–2021) is from the BC Sablefish operating model.

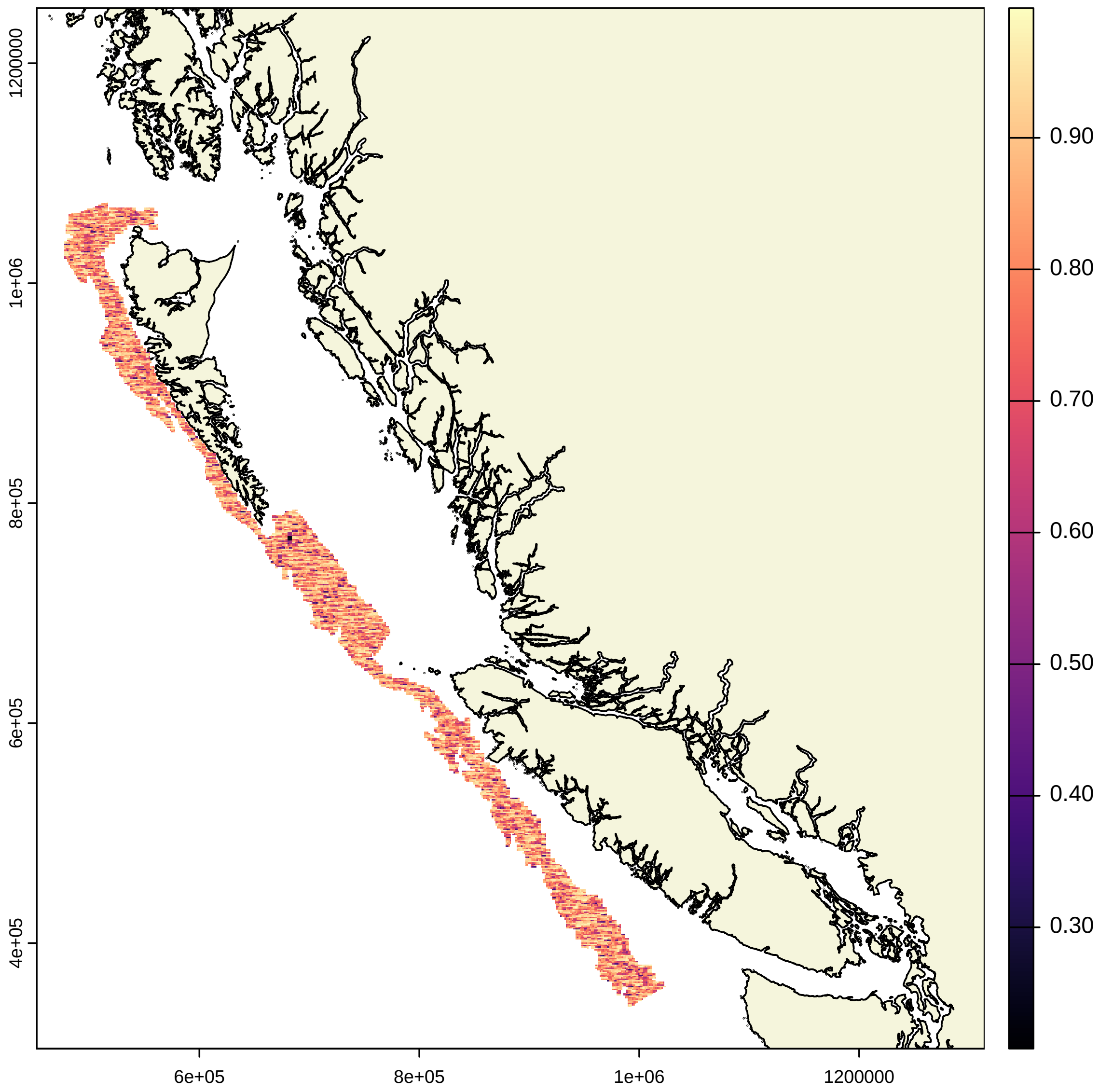


Figure S.9: The 0.025 quantile for estimates of RBS for sponge habitats in 2024 for $i$ = 25 081 grid cells (1 km x 1 km) considered in risk assessment. Coastline data are from Wessel and Smith 1996 and map projection is NAD83 BC Albers (EPSG:3005).

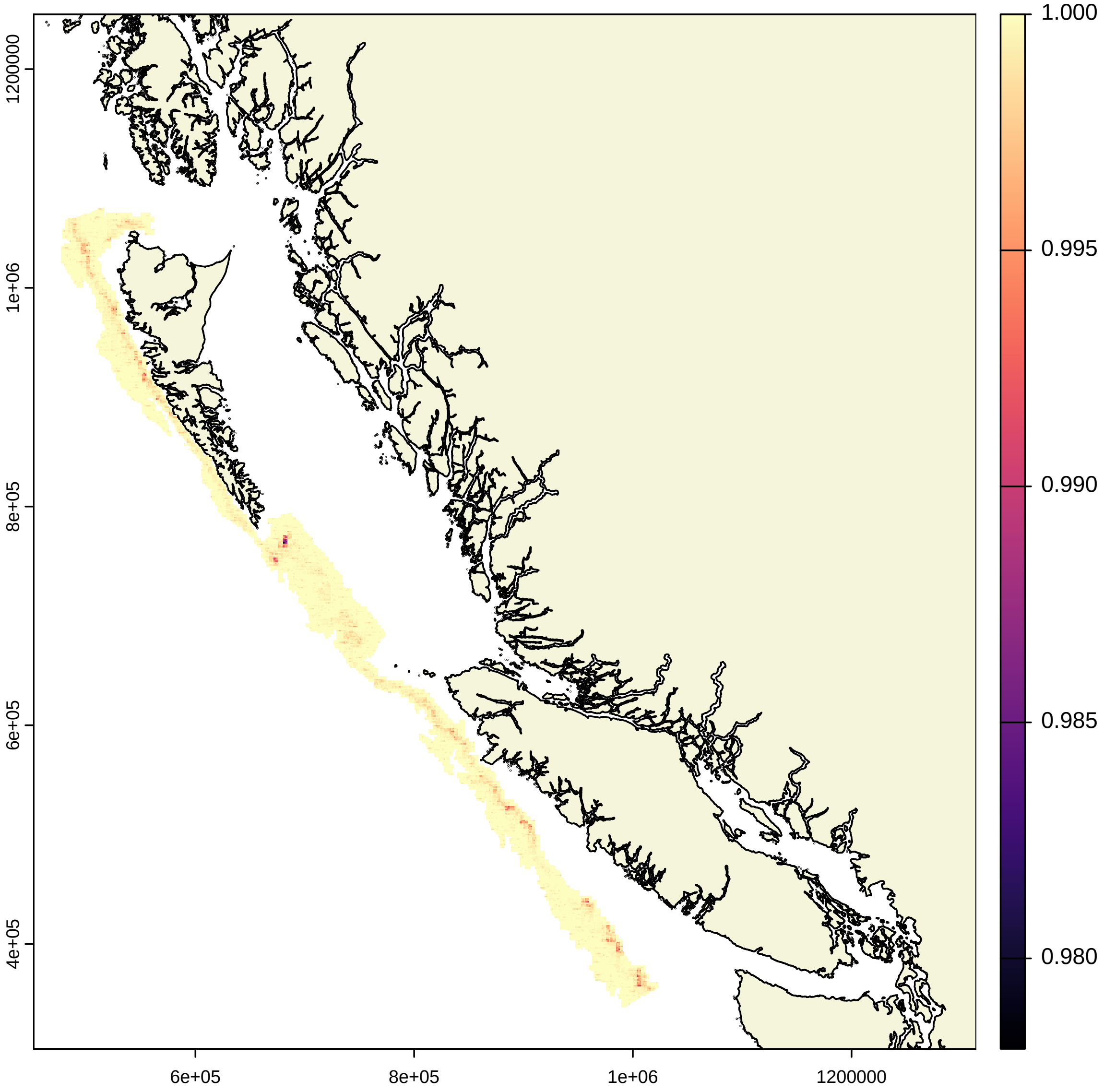


Figure S.10: The 0.975 quantile for estimates of RBS for sponge habitats in 2024 for $i$ = 25 081 grid cells (1 km x 1 km) considered in risk assessment. Coastline data are from Wessel and Smith 1996 and map projection is NAD83 BC Albers (EPSG:3005).

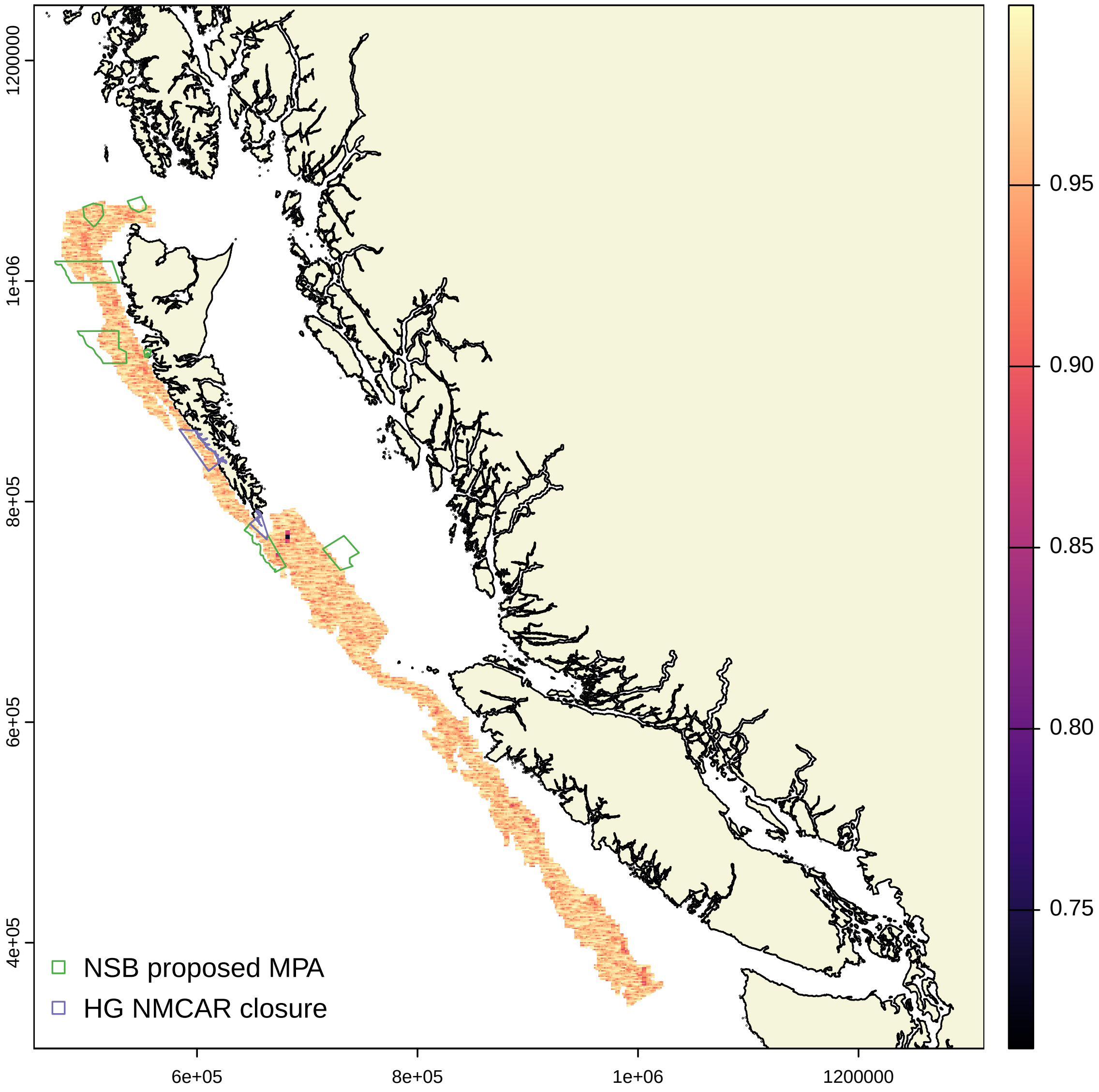


Figure S.11: Mean RBS for sponge habitats in 2024 for $i$ = 25 081 grid cells (1 km x 1 km) examined, including overlap from current and proposed MPAs. Coastline data are from Wessel and Smith 1996 and map projection is NAD83 BC Albers (EPSG:3005).
